# Reduced TBX5 dosage undermines developmental control of atrial cardiomyocyte identity in a model of human atrial disease

**DOI:** 10.1101/2025.08.16.669546

**Authors:** Irfan S. Kathiriya, Kavitha S. Rao, Alexander P. Clark, Kevin M. Hu, Zoe L. Grant, Megan N. Matthews, Zhe Chen, Swetansu K. Hota, Jeffrey J. Saucerman, Benoit G. Bruneau

## Abstract

While atrial septal defects (ASDs) and atrial fibrillation (AF) present differently, there is evidence that they share some genetic basis. Here, we used directed differentiation of human induced pluripotent stem cells into atrial or ventricular cardiomyocytes (CMs) to delineate gene regulatory networks (GRNs) that define each identity. We uncovered accessible chromatin regions, transcription factor motifs and key regulatory nodes specific to, or shared by, both CM types, including the transcription factor *TBX5*, which is linked to genetic susceptibility of ASDs and AF in humans. Complete *TBX5* loss resulted in a near absence of atrial CMs with a concomitant increase in the abundance of other cell types. Reduced dosage of TBX5 in human atrial CMs caused cellular, electrophysiologic and molecular phenotypes consistent with features of atrial CM dysfunction. This included dose-dependent aberrant accessibility of many chromatin regions and perturbation of gene regulatory networks of atrial CM identity. These results suggest that, in addition to stemming from ion channel or extracellular matrix dysfunction, atrial diseases such as ASDs or AF may result from disruptions of atrial CM identity.

## Introduction

Thin-walled atria are distinct from thick-walled ventricles in their developmental origin and function. Atria serve to collect blood in the heart, initiate and propagate an electrical signal for coordinated cardiac contraction, and respond to physiologic changes such as heart failure by secreting hormones. These functions reflect the distinct phenotypes of atrial cells. For example, atrial fibroblasts display a greater response to pathologic stimuli than ventricular fibroblasts (Burstein et al., 2008), and atrial cardiomyocytes (CMs) display distinct electrophysiologic characteristics, calcium-handling and contractile proteins, cell-cell coupling and energy metabolism, which are reflected in human atrial gene expression *in vivo* (Dobrev et al., 2019).

Atrial disease is common and increases cardiovascular and cerebrovascular risk. Developmental defects of the atria include cor triatriatum, atrial aneurysms, patent foramen ovale (PFO) and atrial septal defects (ASDs). ASDs are one of the most common forms of congenital heart defects (CHDs), affecting 25% of the CHD population (Hoffman and Kaplan, 2002). Some large ASDs can lead to heart failure and require surgical repair. With aging, atrial tissue can manifest chaotic electrical activity known as atrial fibrillation (AF). AF is the most common arrhythmia and a leading cause of heart failure, stroke and sudden cardiac death (Kornej et al., 2020). Moreover, there is an increasing prevalence of AF in adults with CHDs, and AF risk is higher in patients with ASDs than in the non-CHD population (Miguel and Ávila, 2021). However, the molecular basis of ASDs and AF remains poorly understood, and new insights are needed to develop new diagnostic and therapeutic approaches.

There is ample evidence that atrial diseases share a genetic basis. ASDs are caused by inherited dominant mutations in genes encoding sarcomere proteins (Ching et al., 2005; Matsson et al., 2008; Postma et al., 2011), chromatin modifying enzymes (Boniel et al., 2021) and transcription factors (Basson et al., 1997; Garg et al., 2003; Kirk et al., 2007; Li et al., 1997; Posch et al., 2010; Schott et al., 1998; Yuan et al., 2013). Atrial fibrillation is associated with inherited mutations or variants in genes encoding potassium channels (Chen et al., 2003; Hong et al., 2005; Olson et al., 2006; Xia et al., 2005; Yang et al., 2004), but also sarcomere proteins (Choi et al., 2018; Gudbjartsson et al., 2016; Lee et al., 2018), and transcription factors (Guo et al., 2016; Ma et al., 2016; Roselli et al., 2018; Xie et al., 2013). Thus, some genes associated with ASDs and AF may have overlapping functions, though the mechanisms underlying or linking the two atrial diseases are unclear.

Heterozygous mutations of the transcription factor *TBX5* cause Holt-Oram syndrome, a developmental disorder that often includes ASDs in humans (Basson 1997,Li 1997) and in mice (Bruneau et al., 2001). Conditional loss of *Tbx5* in the endocardium (Nadeau et al., 2010) or dorsal mesenchymal protrusion (Xie et al., 2012) leads to ASDs, suggesting an essential role for TBX5 in atrial septal development. Post-natal deletion of *Tbx5* in atrial CMs leads to AF via a calcium transport mechanism (Dai et al., 2019; Nadadur et al., 2016), and *Tbx5* was shown to be essential for the maintenance of an atrial gene expression program (Sweat et al., 2023). Some inherited *TBX5* mutations cause both ASDs and AF in the same individuals (Bever et al., 1996; Postma et al., 2008). As TBX5 is broadly associated with atrial gene regulation, ASDs and AF, understanding TBX5 function presents an opportunity to decipher the rules of gene regulation in atrial development and disease.

To model human atrial disease *in vitro*, multiple groups have developed methods that promote atrial CM differentiation of human pluripotent stem cells, garnering important insights into the pathways involved (Cyganek et al., 2018; Devalla et al., 2015; Lee et al., 2017; Yang et al., 2022a; Zhang et al., 2019). These protocols have been used on wild-type cells for cell type-specific drug screening (Devalla et al., 2015), and for inducing AF by electrical stimulation (Goldfracht et al., 2019) or with drugs that increase the risk of AF (Shafaattalab et al., 2019). However, human cellular models of genetic susceptibility to atrial disease have not yet been reported.

Here, we deployed a recently developed induced pluripotent stem cell (iPSC) model of Holt-Oram Syndrome (Kathiriya et al., 2021) to study genetic susceptibility to human atrial diseases. We observed disease-relevant cellular, electrophysiologic and molecular deficits of atrial CMs. The deficits in cell physiology and developmental gene expression were more profound than those previously observed in ventricular CMs lacking one or two copies of *TBX5* (Kathiriya et al., 2021). Notably, atrial CM specification failed in the complete absence of *TBX5* and atrial CM identity was hindered by heterozygous loss of *TBX5*. Furthermore, we discovered gene regulatory networks (GRNs) for atrial CM identity that were sensitive to TBX5 dosage, and observed disruptions to key nodes, including ASD– and AF-risk genes, in atrial disease-relevant networks.

## Results

### Modeling atrial CM identity from human iPSCs delineates chamber-specific cellular attributes

As an initial step to model atrial differentiation and disease, we differentiated wildtype (WT) iPSCs (WTC11 parental line) (Miyaoka et al., 2014) into either atrial or ventricular CMs and validated atrial-specific vs. ventricular-specific CM characteristics. Immunostaining showed ANP, a well-known atrial marker, to be enriched in atrial CMs (Fig. 1A-G). By contrast, MLC2V, a ventricular marker, was nearly absent among atrial CMs compared to ventricular CMs (Fig. 1H-N). Multielectrode array (MEA) assays revealed a faster beat rate among atrial CMs and shorter rate-adjusted local extracellular action potentials (LEAP), consistent with atrial CM electrophysiology (Fig. 1O, P)(Nerbonne and Kass, 2005) and with iPSC-derived atrial CMs derived by another protocol (Cyganek et al., 2018).

**Fig. 1.**
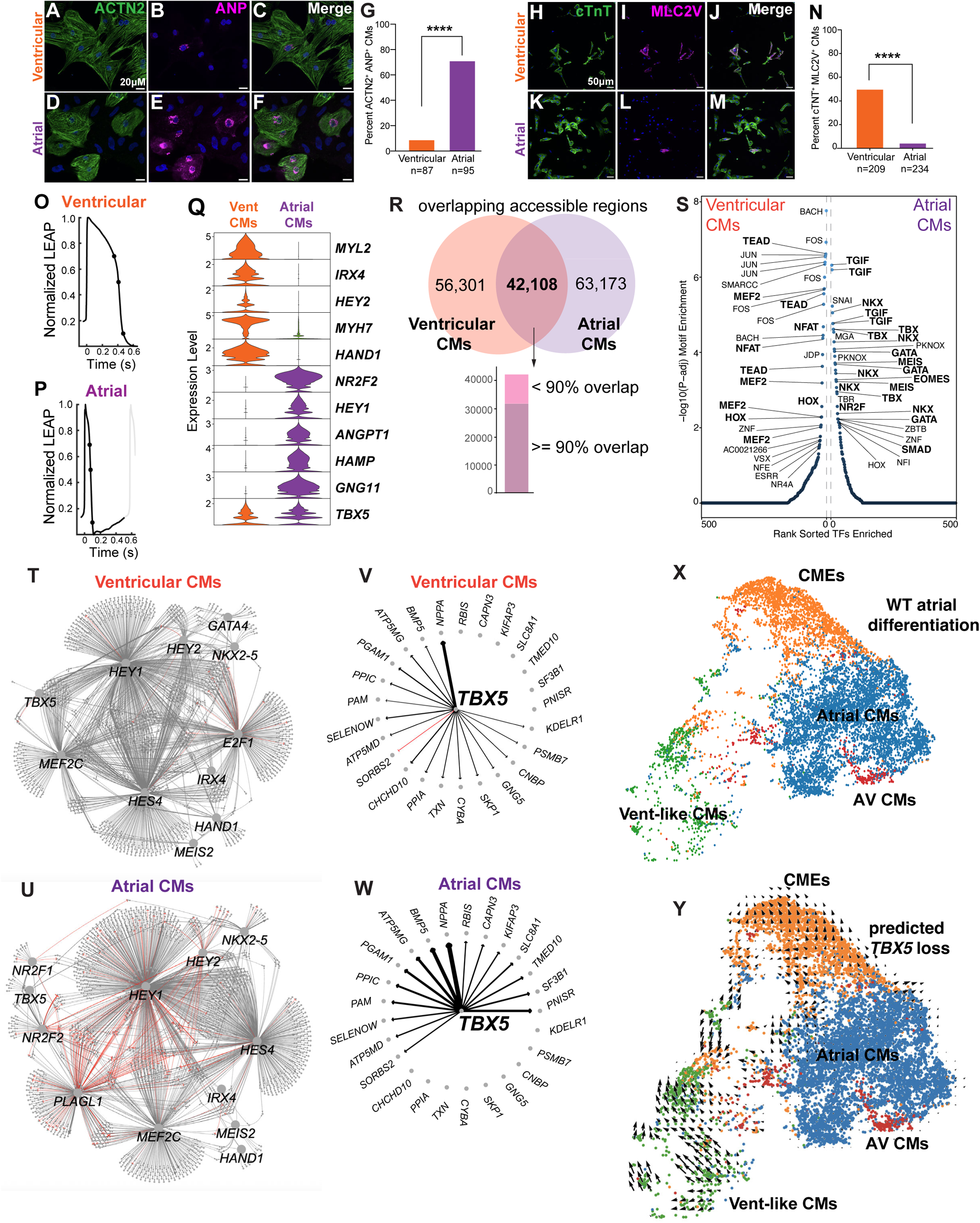
Features of gene expression and chromatin accessibility for regulatory networks of human iPSC-derived atrial cardiomyocytes (CMs) predict TBX5-sensitive atrial CM identity. Directed differentiations of human induced pluripotent stem cells (iPSCs) to ventricular or atrial cells were assessed by immunostaining of (A-G) ANP or (H-N) MLC2V (****p<0.0001 by two-sided Fisher’s exact test). Scale bars denote 20 microns (A-F) and 50 microns (H-M). Cells from two experiments of ventricular or atrial differentiations were analyzed; N depicts number of ACTN2^+^ or cTNT^+^ cells analyzed per group. (O,P) Normalized local extracellular action potential (LEAP) by cell type. Representative traces are shown, based on LEAPs from 6 ventricular or 6 atrial cells from two biological replicates of ventricular or atrial differentiations. (Q) Violin plot of ventricular– or atrial-enriched CM genes. scRNA-seq from two biological replicates of ventricular or atrial differentiations were analyzed. (R) Overlapping chromatin accessible regions. Bar plot shows number of common accessible regions by less than 90% overlap or greater than or equal to 90% overlap. (S) Motifs enriched in differentially accessible regions (DARs) (FDR<0.1). snATAC-seq from two biological replicates of ventricular or atrial differentiations were analyzed in (R, S). Gene regulatory networks (GRNs) of (T) ventricular or (U) atrial CMs. Wheel diagrams for *TBX5* show a shift in significant direct connections between (V) ventricular and (W) atrial CMs. GRNs were derived from two experiments of scRNA-seq or snATAC-seq during ventricular or atrial differentiations. Based on input from the UMAP of atrial CM cell types in (X) wildtype, *in silico* prediction by CellOracle infers the states of individual cells from atrial cardiomyotyces (CMs) to CMs expressing extracellular matrix genes (CMEs) from (Y) complete *TBX5* loss (*TBX5*=0).

We then performed single-cell RNA sequencing (scRNA-seq) on atrial differentiation at day 20 and merged it with previously generated scRNA-seq data of ventricular differentiation at day 23 (Fig. S1A-C) (Kathiriya et al., 2021). Differential expression analysis confirmed that atrial CMs were enriched for atrial genes (e.g. *TBX5*, *NR2F2, HEY1, MYH6*), the right atrial CM gene *HAMP* (Litviňuková et al., 2020; Mir et al., 2023), the left atrial CM gene *PITX2* (Campione et al., 2001; Kitamura et al., 1999; Liu et al., 2002; Schweickert et al., 2000) and the atrial appendage CM gene *NPPA*, but did not express ventricular-specific genes (*MYL2*, *IRX4, HEY2, HAND1, MYH7*) (Fig. 1Q, S1D-E, Table S1).

### Human iPSC-derived atrial CMs display distinct epigenomic features

We next compared accessible chromatin regions between differentiations to ventricular and atrial CMs using single-nucleus ATAC sequencing (snATAC-seq) (Fig. S2A,B, Table S2). We annotated cell type clusters by gene score using ArchR (Granja et al., 2020). Among *TNNT2*-gene score^+^ cells, representing presumptive CMs (Fig. S2B’), *MYL2*-gene score^+^ clusters overlapped with cells from ventricular CM differentiations and *MYH6*-gene score^+^ clusters with cells from atrial CM differentiations (Fig. S2C), as expected. Within a subset of *TNNT2*-gene score^+^ CMs (Fig. S2D,E), we found that accessible chromatin regions were distributed across promoters, introns, exons and distal regions to a similar degree among clusters of ventricular or atrial CMs (Fig. S2F).

By genomic coordinates, about one-quarter of accessible regions (42,108 regions) overlapped between ventricular and atrial CMs at the stages assessed, out of 56,301 ventricular CM-specific and 63,173 atrial CM-specific regions identified (Fig. 1R). Based on fold change differences, we also identified 465 differentially accessible regions (DARs) among ventricular or atrial CMs (Fig. S2G). Thus, although several accessible regions may be common to ventricular and atrial CMs, the degree of accessibility may be different among ventricular or atrial CMs.

Among DARs, transcription factor (TF) motifs enriched among ventricular CMs included MADS-box/MEF2 (Desjardins and Naya, 2016; Medrano and Naya, 2017; Naya et al., 2002), TEAD (Gise et al., 2012; Lin et al., 2016) and NFAT (Bushdid et al., 2003) motifs (Fig. 1S). Atrial CMs were enriched for NKX, NR2F, GATA and TBX or MEIS (Luna-Zurita et al., 2016) motifs (Fig. 1S). Notably, TF motifs of TGIF, a TALE homeodomain protein associated with holoprosencephaly that inhibits retinoic acid signaling (Bartholin et al., 2006) and represses Smad signaling (Nakashima et al., 2018) were observed. However, any potential role for TGIF or TGIF-like factors in atrial development would require experimental evaluation.

### Gene regulatory networks uncover TF nodes of atrial CM identity

Gene regulatory networks (GRNs) reveal a web of “nodes”, TFs that are important for establishment and maintenance of transcriptional programs, and “edges”, which are gene targets connected to their regulatory nodes, thus yielding more information than just changes to gene expression. To uncover GRNs of human ventricular or atrial CM identity, we applied CellOracle, a machine learning-based approach that builds cell type-specific GRNs using both gene expression and chromatin accessibility data (Table S3) (Kamimoto et al., 2023). For each node, we calculated “betweenness centrality”, which measures information flow through each node, and “degree out centrality”, which measures the number of outgoing direct connections to edges for any given node. By these measures, canonical chamber-specific TFs such as *HEY2* and *HAND1* acted as predominant nodes in ventricular CMs, and *NR2F2* and *HEY1* in atrial CMs (Fig. 1T,U, Fig. S3A-F). Notably, *HEY2*, *HES4* and *HEY1* are involved in Notch signaling, suggesting that the Notch pathway is a regulator of both atrial and ventral CM identities. We also identified TFs that are lesser known in the developing heart as nodes, including *E2F1* in ventricular CMs and *NR2F1* and *PLAGL1* in atrial CMs.

Some TFs (e.g. *HEY2*, *HEY1*, *TBX5*) displayed connections in both ventricular and atrial CM GRNs, although the nature of their connections changed depending on ventricular or atrial CM context (Fig. 1T,U, Fig. S3A-F). For example, ventricular-enriched *HEY2* displayed both higher betweenness and higher degree-out centrality in ventricular CMs (Fig. S3E,F), confirming the importance of *HEY2* in a ventricular CM network. In turn, *HEY2* displayed connections with a positive correlation (i.e. directly proportional between the TF node and a target edge), suggesting that *HEY2* positively regulates some ventricular CM genes (Fig. S3G). By contrast, some *HEY2* connections to atrial CM-specific targets reflected negative correlations (i.e. inversely proportional between the TF node and a target edge), suggesting that *HEY2* inhibits some atrial CM genes (Fig. S3H). Like *HEY2* in ventricular CMs, atrial-enriched *HEY1* also displayed both higher betweenness and higher degree-out centrality in atrial CMs (Fig. S3E,F). Interestingly, most *HEY1* connections reflected positive correlations in ventricular CMs (Fig. S3I, while many connections in atrial CMs reflected negative correlations (Fig. S3J), suggesting that *HEY1’*s role is likely more complex than *HEY2’s* in the atrial CM network. *TBX5* showed connections with positive correlations in both atrial and ventricular CMs, some of which were overlapping, implying some shared roles in both ventricular and atrial CM networks (Fig. 1V, W). Although there were more direct connections to target edges in ventricular than atrial CMs, betweenness centrality for *TBX5* was higher in atrial CMs (Fig. S3E), implying a more important role in atrial CMs.

As variants in *TBX5* are associated with ASDs and AF, we hypothesized that *TBX5* plays a central regulatory role in human atrial CMs. To test this hypothesis, we used CellOracle (Kamimoto et al., 2023) to perform *in silico* perturbation of *TBX5* in atrial CM differentiation. In the normal condition, WTC11 cells generated mostly *MYH6^+^* atrial CMs, some *MYH7^+^* ventricular-like CMs and *TNNT2^+^ COL3A1^+^*CMs, which express extracellular matrix genes (CMEs) (Fig. 1X). CellOracle analysis predicted that complete loss of *TBX5* during atrial CM differentiation would hinder specification of atrial CM identity and instead lead toward other CM subtypes, such as CMEs (Fig. 1Y).

### Reduction of TBX5 dosage impairs cellular features of atrial CMs

To confirm the CellOracle prediction, we leveraged a human disease model of *TBX5* haploinsufficiency (Kathiriya et al., 2021). To this end, we differentiated to atrial CMs iPSCs carrying a heterozygous (*TBX5^in/+^*) or homozygous (*TBX5^in/del^*) *TBX5* loss of function (LOF) mutation, or their wildtype parental line (WTC11) (Kathiriya et al., 2021). Western blot confirmed reduced levels of TBX5 in *TBX5^in/+^* cells and absence of TBX5 in *TBX5^in/del^* cells at day 20 of atrial CM differentiation (Fig. S4A, B). While *TBX5^in/+^* cells differentiated to beating atrial CMs, *TBX5^in/del^* cells often had compromised CM differentiation efficiency and rarely produced beating CMs (Fig. 2A). Further, *TBX5^in/del^* atrial CMs could not be recovered beyond day 20, indicating a more severe phenotype in atrial than ventricular CMs (Kathiriya et al., 2021).

**Fig. 2.**
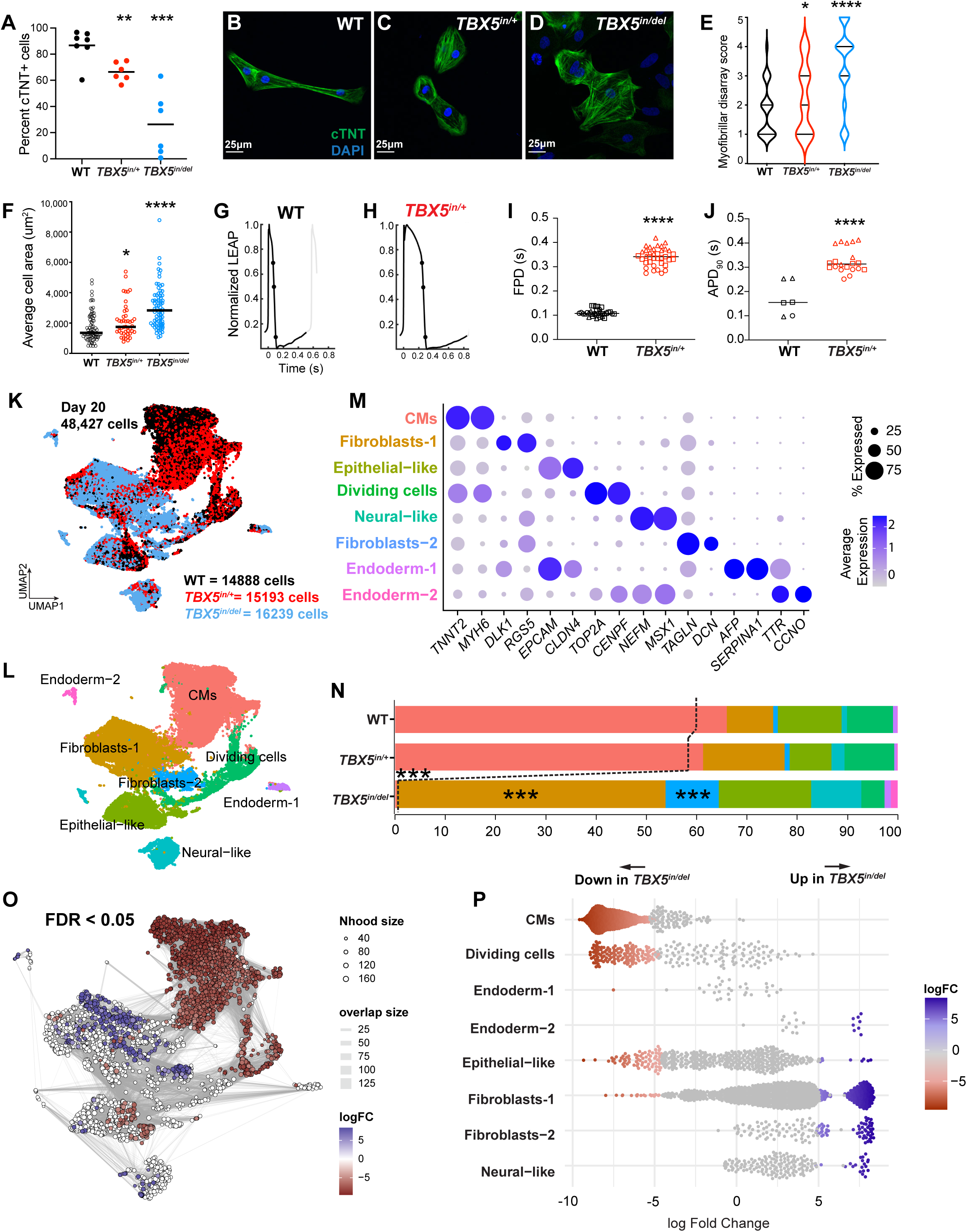
Modeling genetic susceptibility to reduced TBX5 in human iPSC-derived atrial cells reveals atrial CM dysfunction. (A) Differentiation efficiency by flow cytometry of cTNT (**p<0.01, ***p<0.001 by two-sided Student’s T-test). (B-E) cTNT immunostaining shows sarcomeric disarray in *TBX5* mutant atrial CMs (*p<0.05, ****p<0.0001 by two-sided Fisher’s exact test). (F) Average cell area is significantly increased in *TBX5* mutant atrial CMs (*p<0.05, ****p<0.0001 by two-sided Student’s T-test). Cells from two biological replicates were analyzed in (E,F). (G, H) Representative action potential traces by MEA, based on cells from two experiments (N=6 in controls, 20 in *TBX5^in^*^/+^). (I-J) Field potential and normalized local extracellular action potential durations at 90% (APD_90_) are increased in *TBX5* heterozygous mutant atrial CMs (****p<0.0001 by two-sided Welch’s T-test). (K) UMAP displays scRNA-seq data at day 20 by (K) *TBX5* genotype or (L) cell type labels. (M) Dot plot of genes used to define cell types. (N) Distribution of cell types by *TBX5* genotypes shows reduced CMs and increased fibroblasts and epithelial-like cells in *TBX5^in^*^/*del*^ (***adj. p<0.001 by linear regression). *TBX5^in^*^/+^ remains largely similar to WT. (O,P) Milo analysis showed a statistically significant difference in the abundance of CMs, fibroblasts, epithelial and neural-like cells for *TBX5^in/del^*(FDR<0.05). scRNA-seq of two biological replicates by *TBX5* genotype were analyzed.

Furthermore, loss of *TBX5* function led to dosage-sensitive sarcomere disarray and increased cell size at day 20 of differentiation (Fig. 2B-F, Fig. S4C-E), indicating atrial CM dysfunction. Notably, increased cell size in *TBX5^in^*^/+^ atrial CMs was not observed previously in *TBX5^in/+^*ventricular CMs (Kathiriya et al., 2021), again suggesting that atrial CMs are more susceptible than ventricular CMs to reduced TBX5 dosage. Upon replating *TBX5^in/+^* cells for micro-electrode array (MEA), we observed prolonged local extracellular action potentials (LEAPs) with a broad plateau phase (Fig. 2G-J), reminiscent of ventricular-like CM action potentials. This prolonged action potential was consistent with observations from mouse *Tbx5^-/-^* atrial CMs (Yang et al., 2017) and human *TBX5* heterozygous mutant ventricular CMs (Bersell et al., 2023; Kathiriya et al., 2021), albeit using different methods.

### Complete loss of *TBX5* prevents the specification of atrial CMs

To determine the gene expression changes that underlie the phenotypes of *TBX5*-deficient atrial CMs, we performed scRNA-seq on these cells at day 20 of atrial CM differentiation (Fig. 1K, Fig. S4F). A high-level annotation of cell type clusters identified *TNNT2^+^* CMs (WT 60.6% *TNNT2*, *TBX5^in^*^/+^ 59% and *TBX5^in/del^*0.4%), *DLK1^+^* or *DCN^+^* fibroblast-like cells, *EPCAM^+^* epithelial-like cells, *SERPINA1^+^* or *TTR^+^* endodermal population, *TOP2A^+^* dividing cells and *NEFM^+^* neural-like cells (Fig. 2L,M). Using a generalized linear mixed effects model, we tested genotype-specific differences in cluster memberships. Indeed, *TBX5^in/del^* cells were significantly enriched in non-CM clusters of fibroblasts and epithelial-like cells, with a concomitant loss of CMs (Fig. 2N, S4G). As cluster assignments can change with UMAP resolution, we also employed Milo, a program that tests differential abundance by assigning cells to partially overlapping neighborhoods on a k-nearest neighbor graph (Dann et al., 2022). This method also showed a statistically significant change in the abundance of CMs, fibroblasts, epithelial and neural-like cells for *TBX5^in/del^* (Fig. 2O, P). In contrast, cell type abundance was generally similar among *TBX5^in/+^*cells and WT cells at this resolution.

### Reduced TBX5 dosage perturbs the molecular identity of several atrial CM subtypes

Next, we evaluated *TNNT2^+^* clusters and defined CM subtypes (Fig. 3A-C). In WT CMs, we identified CM subtypes that were enriched for *MYH6^+^*expression, such as *PITX2^+^* left atrial (LA)-like CMs, *HAMP^+^*right atrial (RA)-like CMs, *NPPA^+^* atrial appendage-like CMs, *RSPO3^+^* atrioventricular (AV) CMs and a cluster lacking expression of LA– or RA-like markers, which we inferred as atrial precursor-like CMs by pseudotime analysis with Monocle 3 (Cao et al., 2019) (Fig. 3D). Other clusters that had relatively lower *MYH6^+^* expression included *MYH7^+^* ventricular-like CMs expressing ventricular-enriched genes (e.g. *IRX4, HEY2*), *COL3A1*^+^ clusters expressing fibroblast genes (e.g. *COL1A1*, *FN*, *SOX4*, *TGFB2*), resembling CMs expressing extracellular matrix genes (CMEs) in developing mouse hearts (DeLaughter et al., 2016). We also observed a small *HBD^+^* hematopoietic-like cluster.

**Fig. 3.**
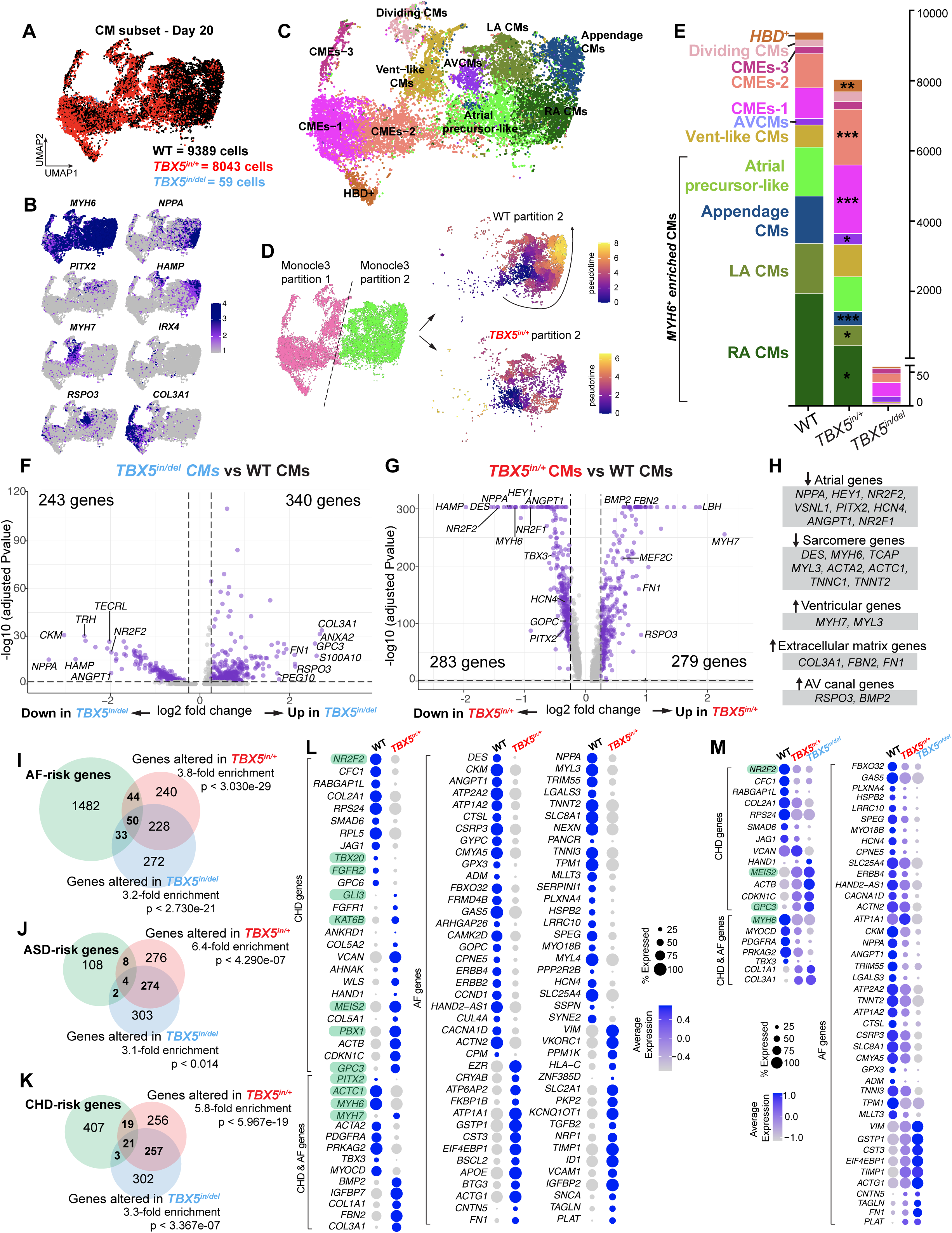
Atrial CM identity by gene expression is vulnerable to reduced TBX5 dosage. UMAP of *TNNT2*+ cells colored by (A) genotype, (B) cell type markers or (C) cell type labels. (D) Monocole 3 analysis shows the predominant pseuodtime paths (i.e., in pseudocolor, represented by the arrows) in two partitions affected in *TBX5^in^*^/*del*^. Specifically, in partition 2, fewer *TBX5^in^*^/*del*^ cells reach the end of the pseudotime path to *NPPA*^+^ CMs. (E) Cell numbers of many atrial CM cell types are significantly reduced in *TBX5* mutants. (*adj. p<0.05 **adj. p<0.01, ***adj. p<0.001 by linear regression). (F,G) Volcano plots show differentially expressed genes in *TBX5* mutants (adj. p-value by Wilcoxan rank sum). (H) Categories of down– and up-regulated genes in *TBX5^in/+^* or *TBX5^in^*^/del^ (adj. p<0.05 by Wilcoxan rank sum). (I-K) AF-, ASD– and CHD-risk genes are enriched among differentially expressed TBX5-sensitive genes (non-adjusted p-value by hypergeometric test). Dot plots of differentially-expressed CHD, ASD (in green) and AF genes in (L) *TBX5^in/+^* or (M) in both *TBX5^in/+^* and *TBX5^in/del^* (adj. p<0.05 by Wilcoxan rank sum). scRNA-seq of two biological replicates by *TBX5* genotype were analyzed.

While similar numbers of total *TNNT2^+^* CMs were observed for WTC11 (9823 cells) and *TBX5^in/+^* (9311 cells), only 59 *TNNT2^+^* CMs were observed from complete loss of *TBX5,* indicating severely impaired atrial CM differentiation, including of most atrial CM subtypes (Fig. 3E, Fig. S5A). *TBX5^in/del^* CMs were devoid of *NPPA*^+^ appendage-like CMs and RA-like CMs but were observed in CME and AV CM clusters. Further, only two atrial precursor-like CMs were observed in *TBX5^in/del^*, indicating an absence of proper atrial CM specification, consistent with *in silico* predictions (Fig. 1X,Y).

Although heterozygous loss of *TBX5* cells were found among RA-like, LA-like and *NPPA^+^* atrial appendage-like CM clusters, *TBX5^in/+^* cells were reduced in numbers, implicating a deficiency of proper atrial CM identity (Fig. 3E, Fig. S5A). Concomitantly, *TBX5^in/+^* cells were enriched for CMEs and AV CMs, consistent with dosage-sensitive expansion of some atrial CM subtypes with loss of others.

In addition, a spectrum of phenotypes from *TBX5* heterozygous mutant cells were observed, with some remaining similar to WT cells, while others were found in a *TBX5* heterozygous-predominant cluster (e.g. CME-1) (Fig. 3A,C,E). Such variability in cellular phenotypes may be of clinical relevance, as many affected members of multi-generational families of Holt-Oram Syndrome, and genetically-determined CHDs in general, express variations in cardiac phenotypes (Basson et al., 1994; Benson et al., 1998; Benson et al., 2003; Garg et al., 2003; Garg et al., 2005).

### ASD– and AF-risk genes are vulnerable to reduced TBX5 dosage

Complete loss of *TBX5* in atrial CMs led to reduced expression of 243 genes and increased expression of 340 genes (Fig. 3F,H, Table S4). Heterozygous loss of *TBX5* resulted in reduced expression of 283 genes and increased expression of 279 genes (Fig. 3G,H, Table S4). Differentially expressed genes were significantly enriched for known AF-, ASD-, or CHD-risk or causative genes (Fig. 3I-K, Table S5). For instance, in *TBX5* heterozygous mutant cells, susceptible genes included 8 ASD (e.g. *TBX20, FGFR2, GLI3, KAT6B, PBX1*), 80 AF (e.g. *DES, CAMK2D, ID1, VCAM1*) and 4 ASD and AF-(e.g. *PITX2, ACTC1, MYH7*) risk genes (Fig. 3L).

Further, dose-dependent genes dysregulated in both *TBX5* mutant genotypes (Fig. 3M) included atrial-specific genes (e.g. *NR2F2*, *HEY1*), the right atrial CM gene *HAMP*, sarcomere genes (e.g. *TNNC1*, *TNNT2*), ECM genes (e.g. *COL3A1*, *FN1*) and AV canal markers (e.g. *RSPO3*, *BMP2*). Many genes could be further categorized as risk genes for ASDs (e.g. *NR2F2, JAG1*) (Ching et al., 2005; McElhinney et al., 2002; Turki et al., 2014), AF (e.g. *NPPA*, *TECRL, TTN)* (Choi et al., 2018; Devalla et al., 2016; Hodgson-Zingman et al., 2008; Lee et al., 2018), or both ASDs and AF (e.g. *MYH6*) (Ching et al., 2005; Lee et al., 2018; Postma et al., 2008; Postma et al., 2011; Roselli et al., 2018; Vad et al., 2020; Yuan et al., 2013).

As expected, many previously characterized TBX5 targets (e.g. *NPPA*, *MYH6, PITX2, DES*) (Bruneau et al., 2001; Garg et al., 2003; Kathiriya et al., 2021; Ouwerkerk et al., 2021) displayed TBX5 dose-dependent dysregulation. We also discovered new putative TBX5 downstream genes, genes such as *ANGPT1, NR2F1* and *MEG3* (Fig. 3F-H). Notably, *ANGPT1*, with known roles in cardiac jelly homeostasis during atrial chamber morphogenesis (Kim et al., 2018), was downregulated in *TBX5* mutant CMs, whereas *MEG3*, a positive regulator of cardiac fibrosis (Piccoli et al., 2017) was downregulated in *TBX5^in^*^/+^ and upregulated in *TBX5^in/del^* CMs. We confirmed altered mRNA expression of *NR2F1, DES*, *ANGPT1* and *MEG3* in *TBX5* mutant atrial CMs by fluorescent *in situ* hybridization (Fig. 4A-M).

**Fig. 4.**
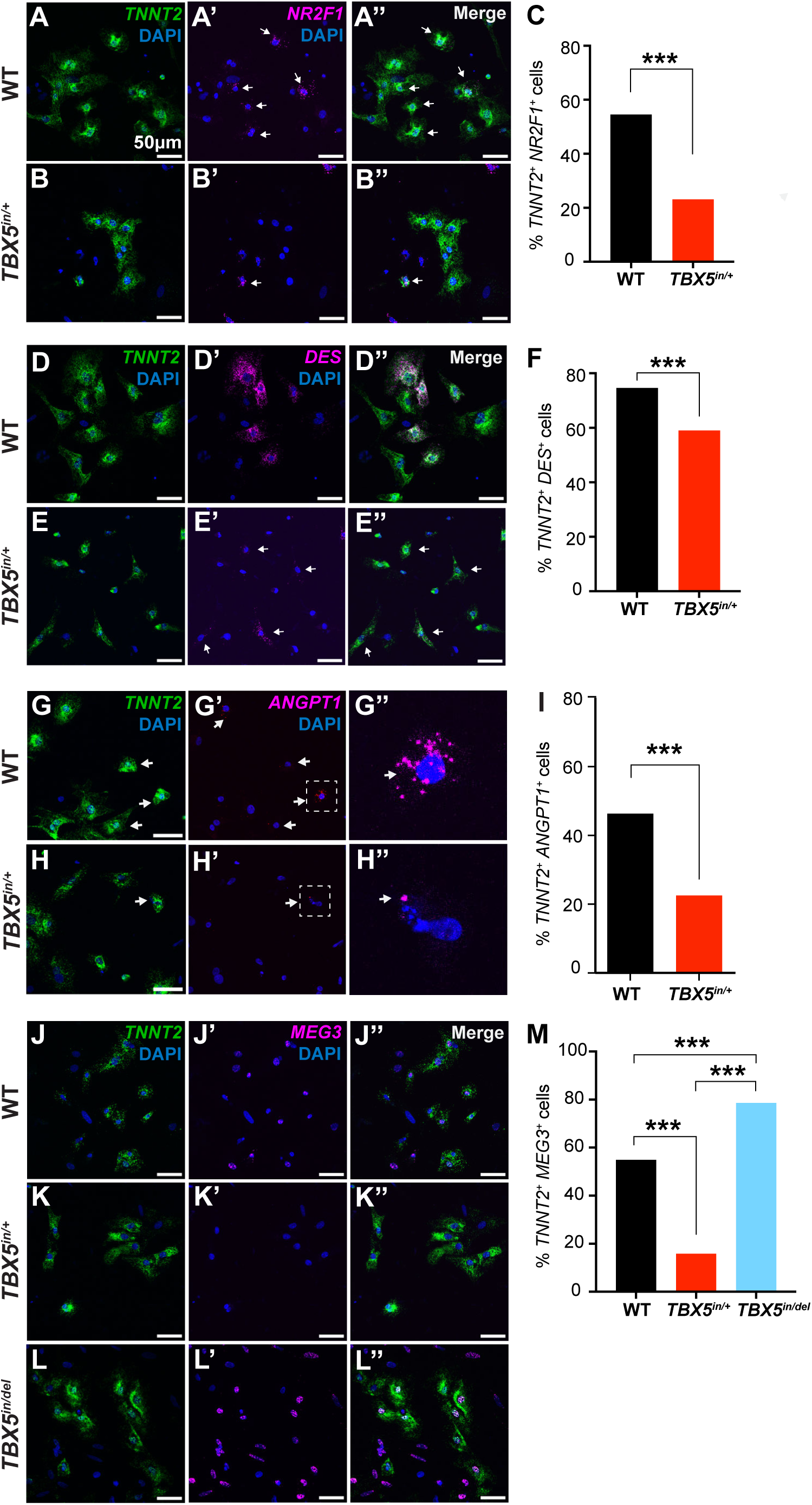
TBX5-dependent genes are dysregulated in human iPSC-derived atrial cells by fluorescent *in situ* hybridization. Gene expression in WT or *TBX5* mutants of *TNNT2* and *NR2F1* (A-C), *DES* (D-F), fibrosis modulator *ANGPT1* (G-I), and cardiac fibrobast-enriched *MEG3* (J-M) (***p<0.001 by two-sided Fisher’s exact test). All scale bars denote 50 microns. Two experiments by *TBX5* genotype were analyzed.

To determine if disturbances from reduced TBX5 are exacerbated later in atrial CM differentiation, we performed scRNA-seq at day 45 in *TBX5^in/+^* atrial cells. Both high-level cell type annotations as well as CM subtypes were similar to those observed at day 20, with the addition of *SHOX2*^+^ pacemaker-like cells (Fig. S6A-G). Fewer *TNNT2*^+^ CMs and more *POSTN*^+^ fibroblasts were observed among *TBX5^in/+^* at day 45 (Fig. S6D). Among CM subtypes at day 45, *HAMP*^+^ RA-like, and *PITX2*^+^ LA-like CMs were reduced, while CME-2s and *MYH7*^+^ ventricular-like CMs were enriched (Fig. S4E-H). These results indicated that *TBX5^in/+^* cells persisted in their failure to differentiate to some atrial CM subtypes, with deviation towards other atrial CM subtypes. Moreover, by day 45, a population of non-atrial CMs, mainly fibroblasts and ventricular-like CMs, expanded. Thus, at day 45, heterozygous loss of *TBX5* appeared analogous, albeit to a lesser degree, to complete loss of *TBX5* at day 20.

Disease genes were also enriched among the 244 upregulated and 154 downregulated genes in *TBX5^in/+^* CMs at day 45 (Fig. S6I). These included 6 ASD-, 72 AF-, and 5 ASD– and AF-risk genes (Fig. S6J-M). A merged analysis of day 20 and day 45 CMs (Fig. S7A,B) revealed genes dysregulated at both timepoints, including 3 ASD-(e.g. *NR2F2, GPC3, MEIS2*), 17 AF-(e.g. *ANGPT1, NPPA, TNNT2, TNNI3*) and 2 ASD and AF-(e.g. *MYH7, ACTC1*) risk genes (Fig. S7C).

### Atrial-enriched TF motifs are sensitive in TBX5-dependent chromatin accessibility

To evaluate whether altered chromatin accessibility might explain disturbances to gene expression by reduced TBX5 dosage in atrial CMs, we performed snATAC-seq analysis of WT and *TBX5* mutant cells at day 20 of atrial CM differentiation (Fig. 5A,B, Fig. S8A). Among *TNNT2* gene score^+^ clusters, cluster C5 was enriched for *TBX5^in/del^* cells, which showed reduced gene scores for *MYH6*, *PITX2, HAMP* and *NPPA,* and increased gene scores for *COL3A1*, indicating disrupted chromatin accessibility of CMs from complete *TBX5* loss. In contrast, cluster C6 was enriched for *TBX5^in^*^/+^ cells and displayed reduced gene scores for *PITX2*, *HAMP* and *NPPA*, consistent with disturbed chromatin accessibility of atrial CMs from heterozygous loss of *TBX5* (Fig. 5C, D, Fig. S8B).

**Fig. 5.**
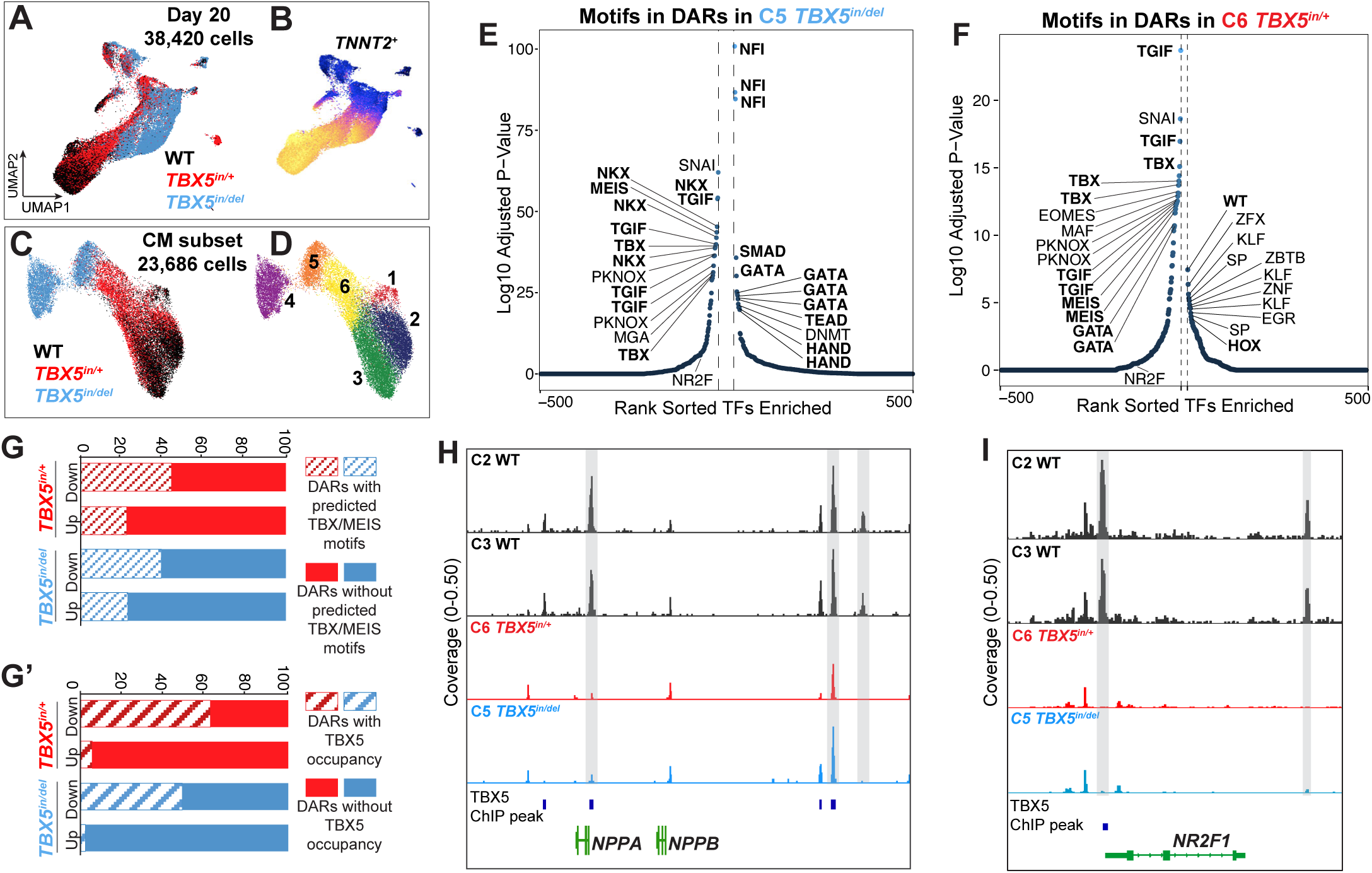
TBX5-sensitive chromatin regions in human iPSC-derived atrial cells show reduced accessibility at sites normally bound by TBX5. UMAP of all atrial cells colored by (A) *TBX5* genotype or (B) *TNNT2*^+^-gene score. UMAP of *TNNT2*^+^-gene score cells colored by (C) *TBX5* genotype or (D) cluster. (E-F) Motifs enriched in differentially accessible regions (DARs) in (E) *TBX5^in^*^/*del*^ or (F) *TBX5^in^*^/+^ (FDR<0.1). DARs with (G) predicted TBX or MEIS motifs or (G’) TBX5 occupancy by TBX5 ChIP-seq. Browser track shows scATAC peaks near (H) *NPPA-NPPB* or (I) *NR2F1*. Gray shading indicates DARs. TBX5 occupancy in WT cells by TBX5 ChIP-seq (Grant et al. 2024). snATAC-seq of two biological replicates by *TBX5* genotype were analyzed.

We analyzed DARs from reduced TBX5 dosage for TF motif loss or enrichment. We found a loss of atrial-enriched NKX, TBX or MEIS and TFIG motifs between WT-enriched and *TBX5^in/del^*-enriched clusters, and similar loss of NKX, TBX or MEIS, GATA, NR2F and TGIF motifs between WT-enriched and *TBX5^in/+^*-enriched clusters (Fig. 5E,F, S8C-E, Table S6). These TF motifs suggested a broadly disturbed deployment of cardiac TFs determining atrial CM identity in *TBX5* mutants. In contrast, TF motifs gained included non-heart associated NFI, ventricular and outflow tract-associated HAND, and BMP-responsive SMAD motifs in *TBX5^in/del^* atrial CM and motifs for the epicardial-associated WT1 in *TBX5^in^*^/+^ atrial CMs (Fig. 5E,F). These changes suggested non-atrial CM TFs had gained accessibility to CM chromatin, consistent with ectopic expression of non-atrial CM genes (Fig 3H).

As TBX and MEIS motifs share sequence similarity (Luna-Zurita et al., 2016), we evaluated DARs with predicted TBX or MEIS motifs. We observed that approximately 40-60% of regions less accessible in *TBX5* mutant CMs contained TBX or MEIS motifs compared to about 20% of regions more accessible in *TBX5* mutant CMs (Fig. 5G), implying that TBX5 may be necessary for open chromatin. Further, to assess if changes in chromatin accessibility could be a direct effect of reduced TBX5 dosage, we overlaid TBX5 occupancy from TBX5 ChIP-seq in WT cells (Grant et al., 2025) on changes to chromatin accessibility in *TBX5* mutants. We found that 50-60% of the regions less accessible in *TBX5* mutants overlapped with normal TBX5-binding regions, whereas less than 5% of the more accessible regions did (Fig. 5G’). This observation suggests that reductions in chromatin accessibility are likely a direct effect of reduced TBX5 binding to its target sites. Evaluation of chromatin accessibility near differentially expressed genes showed correlations between gene expression and chromatin accessibility (Fig S8F-K). Some examples of DARs across genotype-enriched clusters included the proximal promoter of the AF-linked gene *NPPA* (Horsthuis et al., 2008), the nearby *NPPA-NPPB* super enhancer (Man et al., 2021) (Fig. 5H), as well as the promoter of the novel TBX5 atrial CM target *NR2F1* (Fig. 5I).

### Loss of one *TBX5* copy abolishes TBX5 centrality and rewires atrial GRNs

We used CellOracle (Kamimoto et al., 2023) to infer GRNs of atrial CMs by *TBX5* genotype and to understand how reduced TBX5 dosage perturbs network connections for human atrial CM identity (Table S7). Decreasing TBX5 dosage reduced the total number of nodes from 1106 in WT to 1028 in *TBX5^in/^* and 941 in *TBX5^in/del^* (Fig. S9A), with a concomitant increase in genotype-specific edges from 900 in WT to 1436 in *TBX5^in/+^* and 1625 in *TBX5^in/del^* (Fig. S9B). Many network connections displayed shifts from positive correlations to negative correlations (e.g. *NR2F1*, *NKX2-5*, *HEY1, TCF21*) (Fig. 6A-C, Fig. S9C-F). Furthermore, a network comprised of upstream TFs connected to TBX5 or CHD-, ASD– and AF-risk TFs susceptible to reduced TBX5 dosage showed fewer unique edges (61 in WT, 47 in *TBX5^in/+^,* 36 in *TBX5^in/del^*) and total nodes (27 in WT, 25 in *TBX5^in/+^,* 19 in *TBX5^in/del^*) (e.g. ASD-risk *SMC3* lost in *TBX5^in^*^/+^ and *TBX5^in^*^/*del*^, *MEF2A* lost in *TBX5^in^*^/+^) (Fig. 6D-F). These results indicated that these networks are less complex when TBX5 dosage is reduced, implying a loss to a TBX5-dependent atrial CM program.

**Fig. 6.**
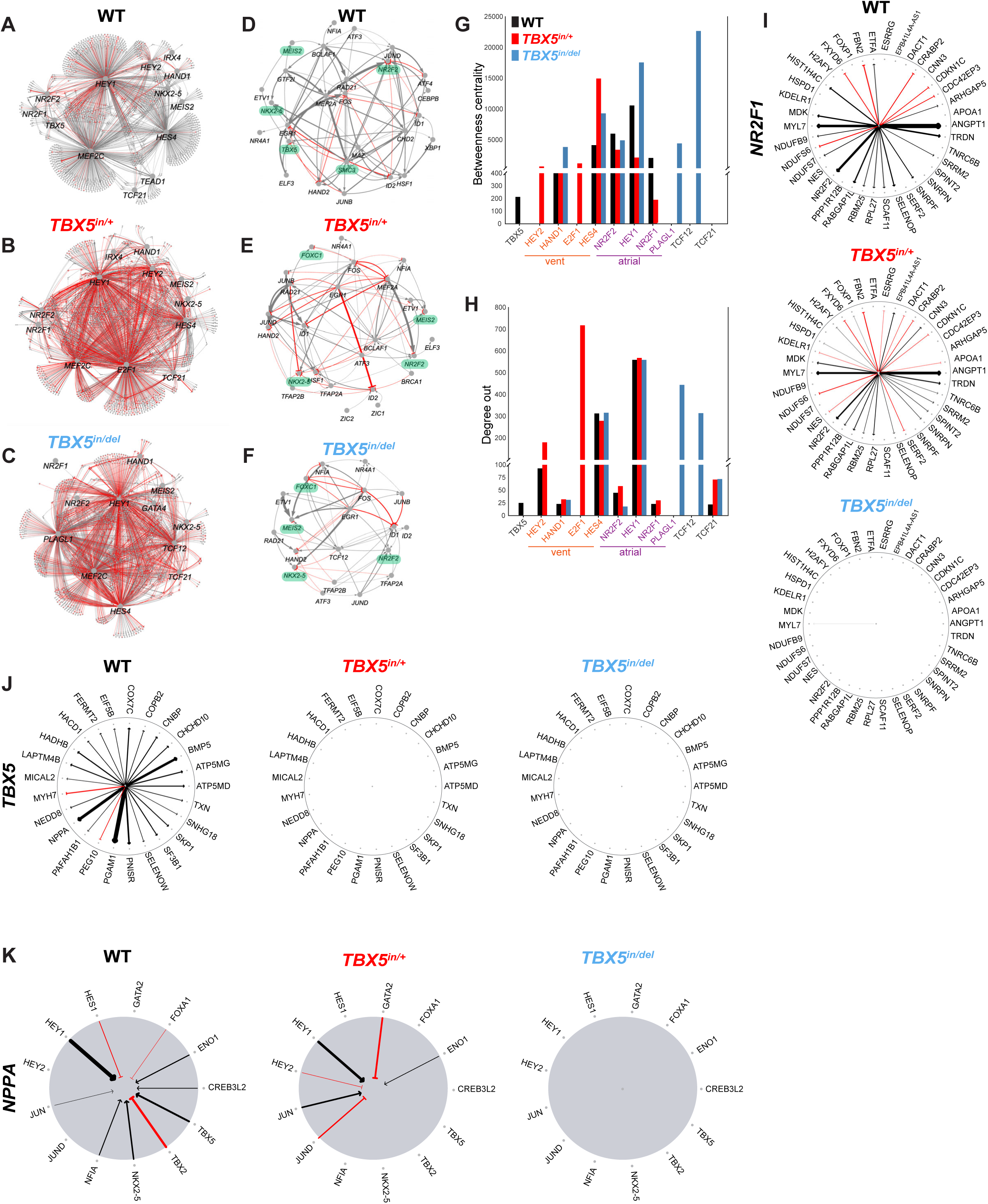
Proper TBX5 dosage controls disease-linked atrial CM gene regulatory networks (GRNs). (A-C) GRNs of atrial CMs with selected TFs by *TBX5* genotype. Grey lines indicate positive correlations. Red lines depict negative correlations. Genes that are edges are unlabeled for simplicity of the diagram. (D-F) Priority networks of atrial CMs shows TF sources to TBX5 or CHD-, ASD-(in green) or AF-risk genes in *TBX5* mutants. (G) Betweenness or (H) degree out centrality of selected TFs in GRNs. Wheel diagrams of (I) *NR2F1* or (J) *TBX5*. Strength of black arrows depicts strength of positive correlation, while strength of red arrows depicts strength of negative correlations. (K) Inverse wheel diagram of *NPPA* shows connections to other TFs that are gained or lost from reduced TBX5 dosage. Two samples of scRNA-seq or snATAC-seq by *TBX5* genotype were analyzed.

Several TFs showed dosage-sensitive reduction in centrality by betweenness or degree out (Fig. 6G,H). *NR2F1* maintained many of its degree-out connections in *TBX5^in^*^/+^ cells, but the strength of correlations, if not their direction, was altered (e.g. AF-risk *ANGPT1*, *MYL7, ASD-risk NR2F2, SELENOP*) (Fig. 6I). Other *NR2F1* connections were lost (e.g. *NDUFB9*, *NDUFS7, ETFA, ARHGAP5, CDC42EP3*) or gained (e.g. *NES*, *PPP1R12B, KDELR, APOA1, CNN3*), consistent with a reconfiguration of *NR2F1* connections in response to reduced TBX5. Correspondingly, the degree-out centrality of *TCF21,* which was low in WTC11 CMs, dramatically increased in *TBX5^in^*^/+^ and *TBX5^in^*^/*del*^ CMs (Fig. S9E), suggesting an ectopic fibroblast-like network in *TBX5* mutants, potentially in conjunction with an increase in connections of the fibroblast-associated *TCF12* (Fig. S9F).

Moreover, *TBX5* lost both its betweenness and degree-out centrality in *TBX5^in/+^* GRNs (Fig. 6G, H). This included loss of all 25 connections, including with the highest correlated gene *PGAM1*, along with *BMP5,* AF-risk *NPPA*, and ASD– and AF-risk *MYH7* (Fig. 6J), consistent with profound disturbances to *TBX5* connections from heterozygous loss of *TBX5* in atrial CMs. Notably, some TBX5 dosage-sensitive edge genes that lost direct connections to TBX5 appeared to form new connections with other transcription factors, including both positive and negative regulators (Fig. 6K, Fig. S10A-C). For example, the TBX5-dependent edge gene *PNISR* gained positive connections to the ASD– and AF-risk TF *NKX2*-*5* and the cardiopulmonary disease-linked TF *TBX4* (Prapa et al., 2022) in *TBX5^in^*^/*del*^ CMs (Fig. S10B). Concurrently, *PNISR* showed reduced strength of a negative connection to the disease-risk TF *TBX2* (Liu et al., 2018) in *TBX5^in/+^,* while gaining a positive connection to *TBX2* in *TBX5^in^*^/*del*^ CMs (Fig. S10B). This suggested some degree of buffering or compensation, albeit incomplete, by other TFs in the setting of reduced TBX5 dosage. In sum, we found that reduced TBX5 dosage results in a reconfiguration of atrial CM networks, including disruptions to connections of ASD– and AF-risk genes.

## DISCUSSION

This study describes a human cellular model of TBX5-associated atrial disease and reveals dosage-sensitive disruptions to atrial CM GRNs. Our findings uncover vulnerabilities in atrial CM identity and provide further evidence that ASDs and AF share a molecular basis. These insights advance our understanding of genetic susceptibility to atrial diseases and provide new potential avenues for therapeutic development.

Using human differentiation systems to CMs, we discern epigenetic differences between closely related cell types, ventricular and atrial CMs. These results provide further support that these approaches may be useful to model *in vitro* cardiac diseases that affect one cell type versus the other. For example, iPSC-derived ventricular CMs have been valuable to model susceptibility to ventricular-specific diseases, such as hypertrophic or dilated cardiomyopathy, ventricular arrhythmias, or congenital heart defect networks (Briganti et al., 2020; Friedman et al., 2024; Hinson et al., 2015; Hnatiuk et al., 2021; Kathiriya et al., 2021; Knollmann, 2013; Lan et al., 2013; Seeger et al., 2019; Sinnecker et al., 2013; Smith et al., 2018; Sun et al., 2012). Similarly, human iPSC-derived atrial CMs can be leveraged for modeling AF, ASDs or other atrial diseases, and to uncover atrial-relevant disturbances to GRNs and cellular physiology. In turn, human cellular models of atrial disease could be used to screen drugs for therapeutic potential, teratogenicity, and atrial arrhythmia risk, and to decipher non-coding variants or oligogenic inheritance of human atrial diseases.

From our data, modeling GRNs of atrial or ventricular CM identity predicted key nodes that could be critical for disrupting atrial GRNs. One of these key nodes is *TBX5*, which we and others have studied (Bersell et al., 2023; Karakikes et al., 2017; Kathiriya et al., 2021). Here, we have modeled reduced dosage, including absolute deficiency of *TBX5 in silico* and *in vitro*, in human atrial cells. Based on changes in gene expression and chromatin accessibility, we have inferred exquisitely sensitive GRNs that are important for atrial function and identity. It is particularly striking that direct connections of multiple key nodes, such as *NR2F1* and ASD-risk *NR2F2*, are affected within these dosage-vulnerable networks even from partial loss of *TBX5*, leading to a cascading effect on susceptible downstream genes related to atrial CM identity. Conversely, the gain of connections of fibroblast-associated *TCF21* and *TCF12* may reflect a change in CM cell identity to CMEs, which express some fibroblast-like genes. In the setting of genetic susceptibility to AF, for example, this perturbation of cell identity may contribute to an atrial fibrosis-like state associated with AF (Platonov, 2017). For some Holt-Oram syndrome patients, we speculate that in atrial tissue, the number of fibroblasts could be increased, while CMEs could be expanded at the expense of other atrial CM subtypes.

Some nodes in the GRNs that define atrial CM identity are associated genetically with AF and/or ASDs, suggesting a common disease network that may underly their pathologies. These disrupted networks point to new TBX5-sensitive candidate genes, such as *NR2F1*, *ANGPT1* and *MEG3*, and their biological processes as possible leads for therapeutic potential. Moreover, reverting dosage-sensitive perturbations to more “normal” GRNs may offer therapeutic opportunities to restore proper atrial identity in patients with genetic or acquired atrial conditions, as demonstrated in another cardiovascular disease context (Theodoris et al., 2020). By combining computational modeling with analysis of this cellular disease model, predictions to enhance TBX5 activity or improve disturbed atrial GRNs could be tested to mitigate atrial disease susceptibility by small molecules or gene editing.

In conclusion, our discovery of a broad disruption to the GRNs underlying atrial identity in the context of genetic susceptibility to atrial disease suggests that some atrial diseases, such as ASDs or AF, may not only arise from disturbances of certain functional proteins, such as ion channels or extracellular matrix components. Rather, some atrial diseases may be caused by a fundamental disruption of atrial CM identity. More generally, we propose that perturbations to cell identity may serve as a basis for genetic susceptibility underlying some human diseases.

## STUDY LIMITATIONS

Several limitations of this study should be considered. Although this human cellular model mimics certain disease phenotypes, ASDs and AF manifest in a 3-dimensional tissue and cannot be fully recapitulated in a 2-dimensional model system. Moreover, certain *in vivo* cell types may be missing or represented poorly *in vitro*. Likewise, contributions from the spatial organization of cells *in vitro* or in heart tissue *in vivo* are not captured. Given that there are several sub-categories of ASDs (e.g. primum ASD, secundum ASD, sinus venosus ASD, or coronary sinus ASD) or AF (e.g. persistent AF, paroxysmal AF, or post-operative AF), it remains to be determined how well this cellular disease model may reflect these conditions. Further, by using scRNA-seq and snATAC-seq separately, we uncovered correlations between gene expression and chromatin accessibility, rather than directly linking gene expression to chromatin accessibility in the same cells. Finally, scRNA-seq data is currently available from LV tissue from a Holt-Oram Syndrome patient (Steimle et al., 2024) but not from atrial tissue. As a result, we cannot determine how many of our findings in our iPSC-derived atrial CM model of Holt-Oram Syndrome will be valid in atrial tissue from some Holt-Oram syndrome patients.

## METHODS

### iPSC culture and atrial cardiomyocyte differentiation

In accordance with the Human Gamete, Embryo and Stem Cell Research (GESCR) Committee, which serves as the Stem Cell Research Oversight Committee at UCSF, human iPSCs (WTC11, *TBX5^in/+^* and *TBX5^in/del^*) (Kathiriya et al., 2021) were maintained in mTESR Plus medium (Stemcell Technologies, 100-0276). For directed differentiations, iPSCs were passaged on to GFR Matrigel-coated 6-well plates until 70% confluency and then differentiated according to manufacturer’s instructions using the Stemdiff Atrial Cardiomyocyte Differentiation Kit (StemCell Technologies, 100-0215). iPSCs were authenticated by genotyping for *TBX5* after each thaw of cells and tested for mycoplasma contamination quarterly.

### Immunostaining

Differentiated cells at day 20 from WTC11, *TBX5^in/+^* and *TBX5^in/del^* iPSCs were dissociated using 0.25% trypsin and quenched with cardiomyocyte maintenance medium containing 10% FBS. Cells were resuspended in cardiomyocyte maintenance medium with 10µM ROCK inhibitor (StemCell Technologies, Y-27632), counted and plated onto growth factor-reduced basement membrane matrix (GFR) Matrigel (Corning, 356231) coated-chambered slides (Ibidi, 80381). Medium was replaced after 24 hours with fresh cardiomyocyte maintenance medium. On the following day, cells were fixed in 4% formaldehyde for 15 minutes and washed with PBS. Cells were then incubated in blocking buffer containing 5% goat serum and 0.1% Triton X-100 in PBS for 1 hour at room temperature. Primary antibodies (alpha actinin, Sigma-Aldrich Cat# A7811, RRID:AB_476766, 1:250 (King et al., 2011); MLC2V, Abcam Cat# ab79935, RRID:AB_1952220, 1:200 (Yang et al., 2022a); ANP, GeneTex, GTX109255, 1:250 (Nandi et al., 2021); cTNT, Thermo, Cat#MS-295-P, RRID:AB_61808, 1:500 (Kathiriya et al., 2021)) in blocking buffer were incubated with cells for 2 hours at room temperature. Following washes in blocking buffer, cells were incubated in secondary antibodies at 1:200 dilution (goat anti mouse Alexa Fluor 488, ThermoFisher, A11029, goat anti mouse Alexa Fluor 594, ThermoFisher, A11005). Cells were then washed with blocking buffer and stained with DAPI at a 1:1000 dilution for 2 minutes. Images were acquired using an Olympus FV3000RS and processed with ImageJ. Evaluations of sarcomere disarray or cell size was performed as described previously (Kathiriya et al., 2021) at day 20.

### Replating for multi-electrode array (MEA)

iPSC-cardiomyocytes were harvested as described above. An 8uL droplet of GFR Matrigel in KO DMEM was placed in the center of each well of a BioCircuit MEA Plate (Axion Biosystems, M384-BIO-24) and incubated for 2 hrs at 37C. GFR was gently removed and approximately 50,000 cells in 8-10ul droplets was placed in the center of each well and incubated for 1 hr at 37C. After incubation, additional StemDiff maintenance media with ROCKi was added to each well and incubated for 24 hrs at 37C. Subsequent media changes were performed every other day with StemDiff maintenance media. MEA recordings were acquired after at least 7 days of maintenance.

### MEA Electrophysiology

Electrophysiological data was recorded on the Maestro Edge Micro Electrode Array device (Axion Biosystems). Live recordings of iPSC-cardiomyocyte field potentials and local extracellular action potentials (LEAPs) were acquired using the Cardiac module in the AxIS Navigator Tool. Representative field potential and LEAP tracings from recordings were acquired using the Axion Biosystems Cardiac Analysis Tool. Data collection and analysis was performed using the MEA software (Axion Biosystems) according to manufacturer instructions.

### Fluorescent *in-situ* hybridization

iPSC-derived cardiomyocytes were harvested as described above. Approximately 100,000 cells were then plated on to 3-well chambered slides with removable silicone (Ibidi 80381). Cells were then fixed with 10% neutral buffered formalin for 30 minutes at room temperature, followed by washes with 1X PBS. Serial dehydration with 50%, 70% and 100% ethanol was performed, and slides were stored in 100% ethanol at –20C. On the day of staining, cells were serially rehydrated and pretreated with hydrogen peroxide for 10 minutes and protease III for 10 minutes at a 1:5 dilution ratio. RNAscope Multiplex Fluorescent Reagent Kit v2 (Advanced Cell Diagnostics, 323100) was used to detect *TNNT2* (518991), *ANGPT1* (482901-C2), *DES* (403041-C2), *MEG3* (400821) and *NR2F1* (583751). Finally, silicone chambers were removed from slides, and a coverslip was mounted using Prolong Gold anti-fade mountant. Slides were imaged at 40X magnification on an Olympus FV3000RS confocal microscope and analyzed using FIJI.

### Harvesting cells for single-cell RNA and ATAC sequencing

Human iPSCs from WTC11, *TBX5^in/+^* and *TBX5^in/del^* were differentiated using the Stemdiff Atrial Cardiomyocyte Differentiation Kit (Stemcell Technologies 100-0215) or the Stemdiff Ventricular Cardiomyocyte Differentiation Kit (Stemcell Technologies 05010). For atrial differentiations, two biological replicates of cells were harvested from all three genotypes at day 20 for scRNA-seq (20,000 cells) or snATAC-seq (10,000 cells), or from WTC11 and *TBX5^in/+^* at day 45 for scRNA-seq (20,000 cells), in separate batches according to manufacturer instructions (10X Genomics). For ventricular differentiations, two biological replicates of 10,000 cells were harvested at day 23 for snATAC-sequencing. Cells were dissociated using 0.25% trypsin, quenched with cardiomyocyte maintenance medium containing 10% FBS and then counted. Cell suspensions were processed for single-cell RNA sequencing using Chromium Next GEM Single Cell 3ʹ GEM, Library & Gel Bead Kit v3.1 (10X Genomics). For single nuclei ATAC-sequencing, nuclei were harvested and processed according to manufacturer’s instructions using the Chromium Next GEM Single Cell ATAC Library & Gel Bead Kit v1.1 (10X Genomics). Libraries were sequenced on an Illumina Novaseq6000 S4 or Novaseq X.

### Analysis of single-cell RNA sequencing data

Fastq files were processed using Cellranger count (v6.1.1). Counts from samples from a single timepoint were aggregated using Cellranger aggr without read-depth normalization. Seurat (v4.4) (Hao et al., 2021) was used to analyze the data. Cells with low UMI count plus a high percentage of mitochondrial reads were considered dying and removed. Potential doublets with aberrantly high gene counts were also removed. In the day 20 dataset, cells with UMI counts between 4000-100,000 and gene counts between 500-10,000 were retained. In the day 45 dataset, cells with UMI counts between 4000-50,000 and gene counts between 500-8,000 were retained. Normalization was performed using SCTransform and batch effects were corrected using the SeuratWrapper function RunHarmony. After annotating major cell types, *TNNT2^+^* clusters were subset and further analyzed. Differentially expressed genes with an adjusted p<0.05 from Wilcoxon rank-sum test were reported.

Metrics for cell quality control. After excluding cells that did not pass these quality control metrics, we found that one cluster, at both days 20 and 45, was marked by CM genes (*TTN, TECRL, MYH7B)*, as well as by many mitochondrial genes (*SLC25A37, MT-ND1, MT-ND4, MT-ND2, MT-ATP6, MT-ND3, MT-CYB, MT-ND6, MT-ND5, MT-CO2, MT-CO3, MT-CO1, MT-ND4L*). As this could be an indicator of CMs undergoing apoptosis, we excluded this cluster from downstream analysis.

In Figure 1, single cell RNA sequencing data from WT atrial differentiation was merged and analyzed with previously generated data from WT ventricular differentiation (Kathiriya et al., 2021). Differentially expressed genes were obtained by comparing atrial CM clusters with ventricular CM clusters. In Figure 3, WT cells from clusters highly enriched for *MYH6* were compared to *TBX5^in/+^* or *TBX5^in/del^*cells from other clusters with *MYH6*, similar to the Monocle partitions in Figure 3D.

Volcano plots were generated using the EnhancedVolcano package. Differential genes were intersected with curated lists containing CHD-associated genes, atrial fibrillation-risk genes and genes associated with atrial septal defects. CHD-associated gene list includes our previously curated list from Kathiriya et al., 2021, and genes from the CHDgene database (http://chdgene.victorchang.edu.au/) (Yang et al., 2022b). Atrial fibrillation-risk genes were gathered from the Harmonizome database (https://maayanlab.cloud/Harmonizome/), and (Ouwerkerk et al., 2019; Ouwerkerk et al., 2020; Roselli et al., 2018) (Table S5). The MiloR package (v2.0.0) (Dann et al., 2022) was used for differential cell type abundance testing. The precomputed KNN graph from Seurat was imported using the buildFromAdjacency function. The number of nearest neighbors k was set to 40. Differential abundance results were visualized by setting alpha = 0.05. Monocle 3 (v1.3.7) (Cao et al., 2019) was used for pseudotime analysis. A separate Monocle 3 object was made for WTC11 or *TBX5^in/+^* cells, and a pseudotime trajectory was generated for cells in partition 2. Cells in the atrial precursor-like clusters were chosen as root cells for each Monocle 3 partition using the “order_cells” function. Pseudotime was visualized on the UMAP using the “plot_cells” function.

### Cell abundance testing

A Generalized Linear Mixed Effects Model (GLMM) was implemented in the lme4 (version 1.1-37) R package (Bates et al., 2015) and was used to estimate the associations between cell type-annotations and *Tbx5* genotype (WT, Het or Hom). These models were run separately for each cluster of cells identified within the full data or within subtypes of CMs. The family argument was set to the binomial probability distribution for the GLMM model fit. Cluster membership of cells by sample was modeled as a response variable by a two-dimensional vector representing the number of cells from the given sample belonging to and not belonging to the cluster under consideration. The corresponding sample ID from which the cell was derived was the random effect variable, and the genotype was included as the fixed variable. The resulting P values for the significance estimated log odds ratios were adjusted for multiple testing using the Benjamini–Hochberg method (Benjamini and Hochberg, 1995).

### Analysis of single-cell ATAC sequencing data

Fastq files were processed using Cellranger ATAC count (v2.0). Data was analyzed by following the tutorial for ArchR (v1.0.2) (Granja et al., 2021). Cells with TSS enrichment >= 6 and nFrags >= 1500 were retained. Pseudobulk replicates for each genotype in each cluster were generated with a minimum of 500 and maximum of 1000 cells for each replicate in a sample-aware manner. Peaks were called with MACS2 by setting reproducibility to 2. Volcano plots show differential peaks with FDR < 0.1 and log2FC >0.25. Motif annotations were added from CIS-BP repository using the addMotifAnnotations function and motif enrichment in differential peaks was performed using the peakAnnoEnrichment function with FDR <0.1 and log2FC >0.25. Overlapping ATAC (this manuscript) and TBX5 ChIP peaks (Grant et al., 2025) were attained using findOverlaps from the Genomic ranges package. Browser tracks were generated using IGV Desktop application. In Figure 1, differentially accessible regions were reported between atrial CM clusters and ventricular CM clusters. In Figure 5, differentially accessible regions were reported between WT cells in WT-enriched (C2+C3) clusters and *TBX5^in/+^* cells in *TBX5^in/+^-*enriched (C6) clusters or *TBX5^in/del^* cells in *TBX5^in/del^*-enriched (C5) clusters.

### Gene regulatory network analysis

Gene regulatory network analysis was performed using the CellOracle (v0.18.0) package (Kamimoto et al., 2023). Default parameters were used according to the documentation, unless specified otherwise. For wildtype atrial CM– and ventricular CM-specific GRNs, base GRNs were constructed using scATAC-seq data at day 20 from WTC11 atrial CM and ventricular CM subsets, respectively (Table S3). Cell type-specific GRNs were then constructed by setting alpha = 1000. For *TBX5* genotype-specific GRNs, base GRNs were constructed using scATAC-seq data at day 20 from WTC11 cells (Table S7). The apparent upregulation of *TBX5* mRNA presumably from non-mutated exons (Kathiriya 2021) in *TBX5* heterozygous mutants was manually corrected by halving the raw counts in *Tbx5^in/+^* and setting the raw counts to zero in *Tbx5 ^in/del^* cells, consistent with Western blot data (Fig S1A, B). Cell type-specific GRNs were then constructed by setting alpha = 1000.

Inferred GRNs were filtered to include only the 12,000 edges with largest absolute coefficient values. Centrality calculations for the 12,000 edge networks were calculated using CellOracle’s links.get_network_score() function. PyVis (Perrone et al., 2020) was used to visualize subnetworks. The large nodes in these networks were manually selected TFs and small nodes are the direct targets in the 12,000-edge networks. The edge color was selected to indicate whether a TF activated (grey) or inhibited (red) a target, and edge thickness was scaled based on absolute coefficient magnitude. For networks in (Fig 1T, U), we chose canonical atrial or ventricular TFs, along with some novel TFs – *TBX5, NR2F2*, *MEF2C*, *NKX2*-*5*, *MEIS2*, *GATA4*, *HAND1*, *IRX4*, *HEY2*, *PLAGL1*, *NR2F1*, *HES4*, *E2F1*, and *HEY1*. For networks in Fig 6A-C, the manually selected TFs included *TBX5*, *NR2F2*, *MEF2C, NKX2-5, MEIS2, GATA4, HAND1, IRX4, HEY2, PLAGL1, NR2F1, HES4, E2F1, HEY1, TCF21, TEAD1*, and *TCF12.* For Fig 6D-F, we identified TFs that had source or target connections to TBX5 (Table S7) and prioritized ones that displayed dose-dependent change in centrality by *TBX5* genotype and are CHD– or AF-risk genes. These networks exclude the TFs and only show direct connections between prioritized genes. Wheel diagrams that display direct interactions between TFs and their targets were developed using Python’s NetworkX package (Hagberg et al., 2008), with a circular layout and edge widths scaled by coefficient values.

## ACKNOWLEDGEMENTS

We thank members of the Bruneau and Kathiriya labs for discussion and comments, and of the Gladstone Stem Cell Core, Histology and Microscopy, and Flow Cytometry Core and the UCSF Center for Advanced Technologies for their expert assistance. We thank Reuben Thomas of the Gladstone Bioinformatics Core for valuable feedback on statistical analyses, and Francoise Chanut of Gladstone Communications for editorial assistance. The Gladstone Flow Cytometry Core is funded by NIH S10 RR028962, the James B. Pendleton Charitable Trust and DARPA. Sequencing was performed at the UCSF CAT, supported by UCSF PBBR, RRP IMIA, and NIH 1S10OD028511-01 grants.

## DATA AND RESOURCE AVAILABILITY

Cell lines are available under a material transfer agreement. All relevant data is available on NCBI GEO (Accession GSE285168 and GSE285169) and details of the code are available at GitHub (apclarkva/kathiriya-rao-2025).

**Fig. S1.**
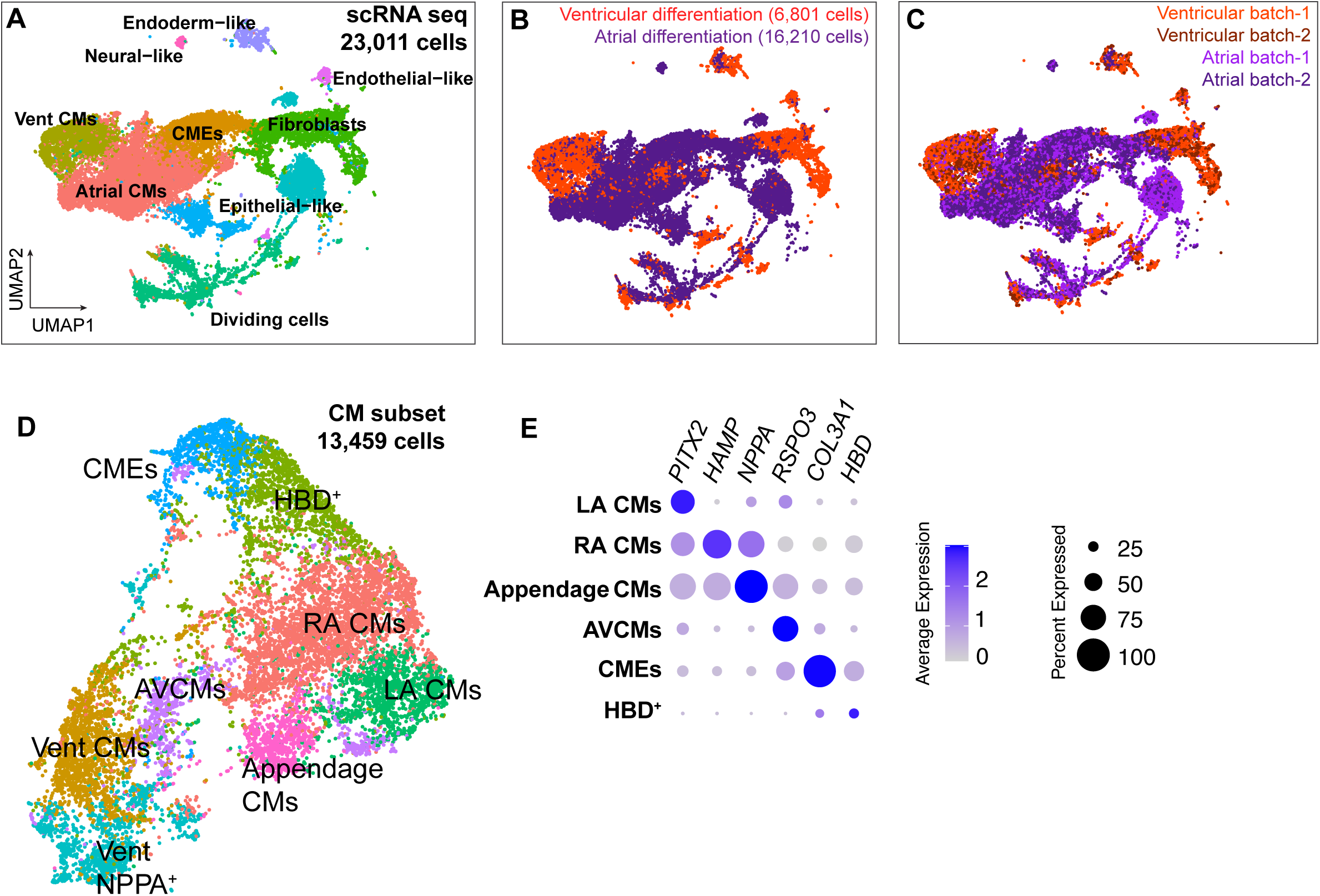
Gene expression analyses in human iPSC-derived ventricular or atrial CMs. UMAPs of scRNA-seq annotated by (A) cell types, (B) differentiation to ventricular or atrial cells, or (C) experimental batch. (D,E) UMAP of CM subset by Louvain cell clustering for subtypes defined by their marker genes. scRNA-seq from two biological replicates of ventricular or atrial differentiations were analyzed.

**Fig. S2.**
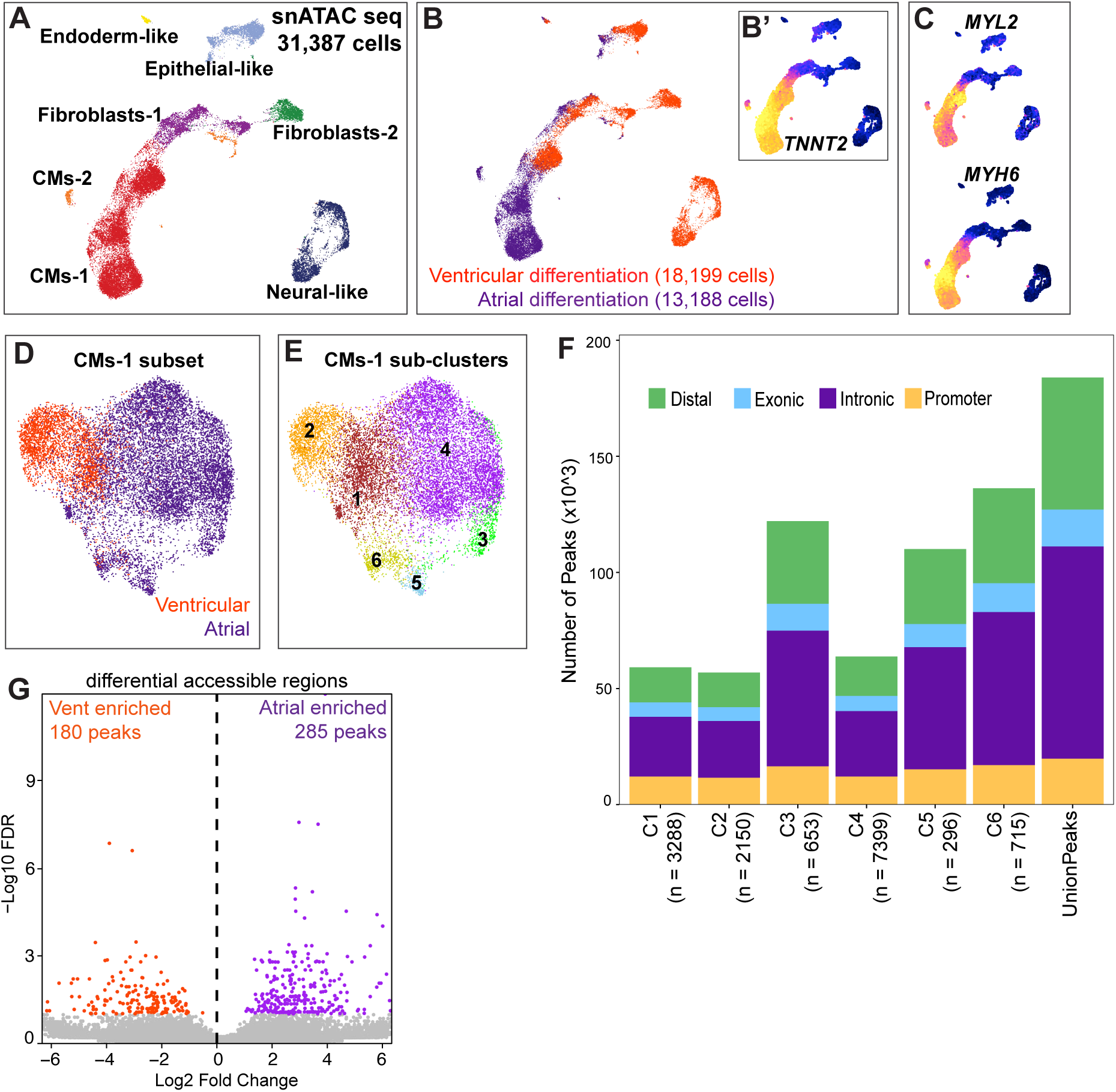
Chromatin accessibility analyses of human iPSC-derived ventricular or atrial CMs. (A) UMAP of snATAC-seq by inferred cell type from gene score, or by (B) differentiation. UMAPs of gene scores from snATAC-seq for (B’) *TNNT2*, (C) *MYL2* (ventricular CM-enriched) or *MYH6* (atrial CM-enriched). UMAP of snATAC-seq subset of *TNNT2*^+^-gene score^+^ cells inferred as CMs (D) by ventricular or atrial differentiation or (E) by Louvain cell clustering. (F) Bar plot of snATAC-seq peak distribution in the genome by cell cluster. (G) Volcano plot of differential accessible regions enriched in ventricular or atrial CMs (FDR<0.1). snATAC-seq of two biological replicates from ventricular or atrial differentiations were analyzed.

**Fig. S3.**
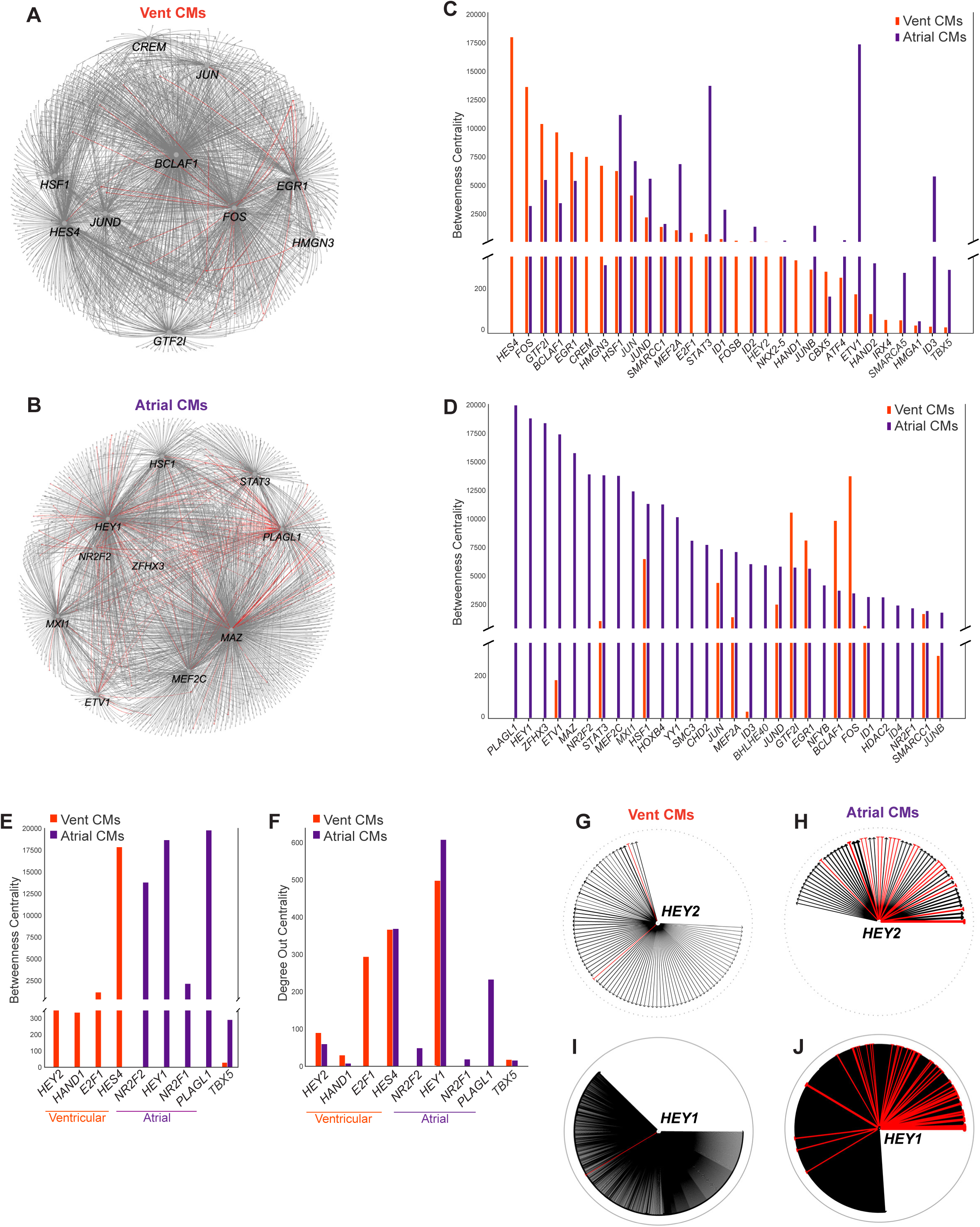
Gene regulatory network (GRN) analyses of human iPSC-derived ventricular or atrial CMs. GRNs display connections of top 10 TFs in (A) ventricular or (B) atrial CMs. Bar plots show top 30 TFs ranked by betweenness centrality scores in (C) ventricular CMs or (D) atrial CMs. Bar plots of selected TFs with known roles in ventricular or atrial CMs show (E) betweenness or (F) degree out centrality. Wheel diagrams of *HEY2* (G, H) or *HEY1* (I, J) display direct connections in ventricular or atrial CMs. GRNs were derived from scRNA-seq or snATAC-seq of two biological replicates during ventricular or atrial differentiations.

**Fig. S4.**
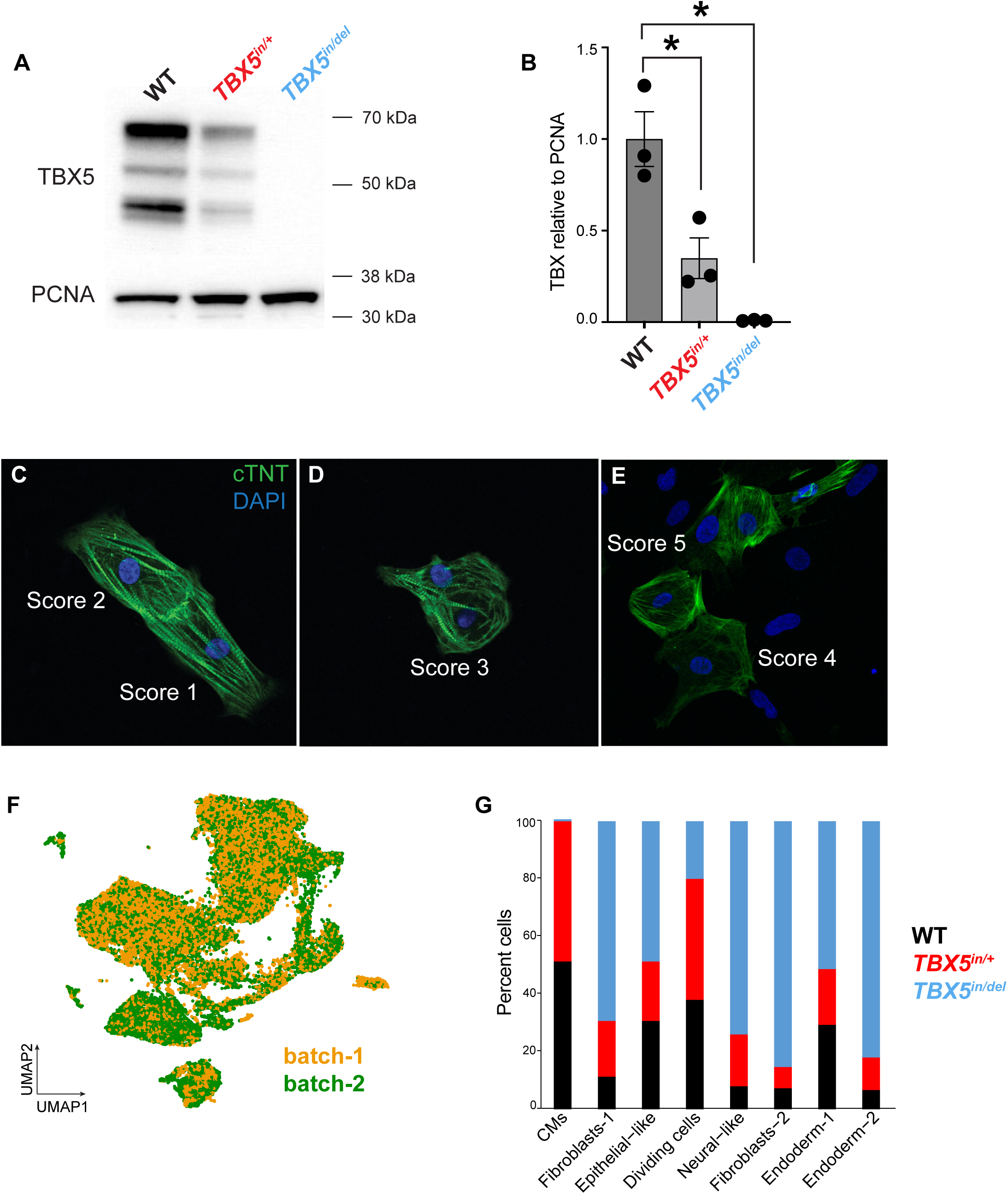
Analysis of a human cellular model of atrial disease. (A) Western blot and (B) quantification of TBX5 by *TBX5* genotype (*p<0.05 by two-sided Welch’s T-test). Representative Western blot is based on three experiments. (C-E) Example images of atrial CMs by sarcomere disarray scores. (F) UMAP of scRNA-seq at day 20 of atrial differentiation by experimental batch. (G) Bar plot shows cell types at day 20 by *TBX5* genotypes. scRNA-seq of two biological replicates by *TBX5* genotype were analyzed.

**Fig. S5.**
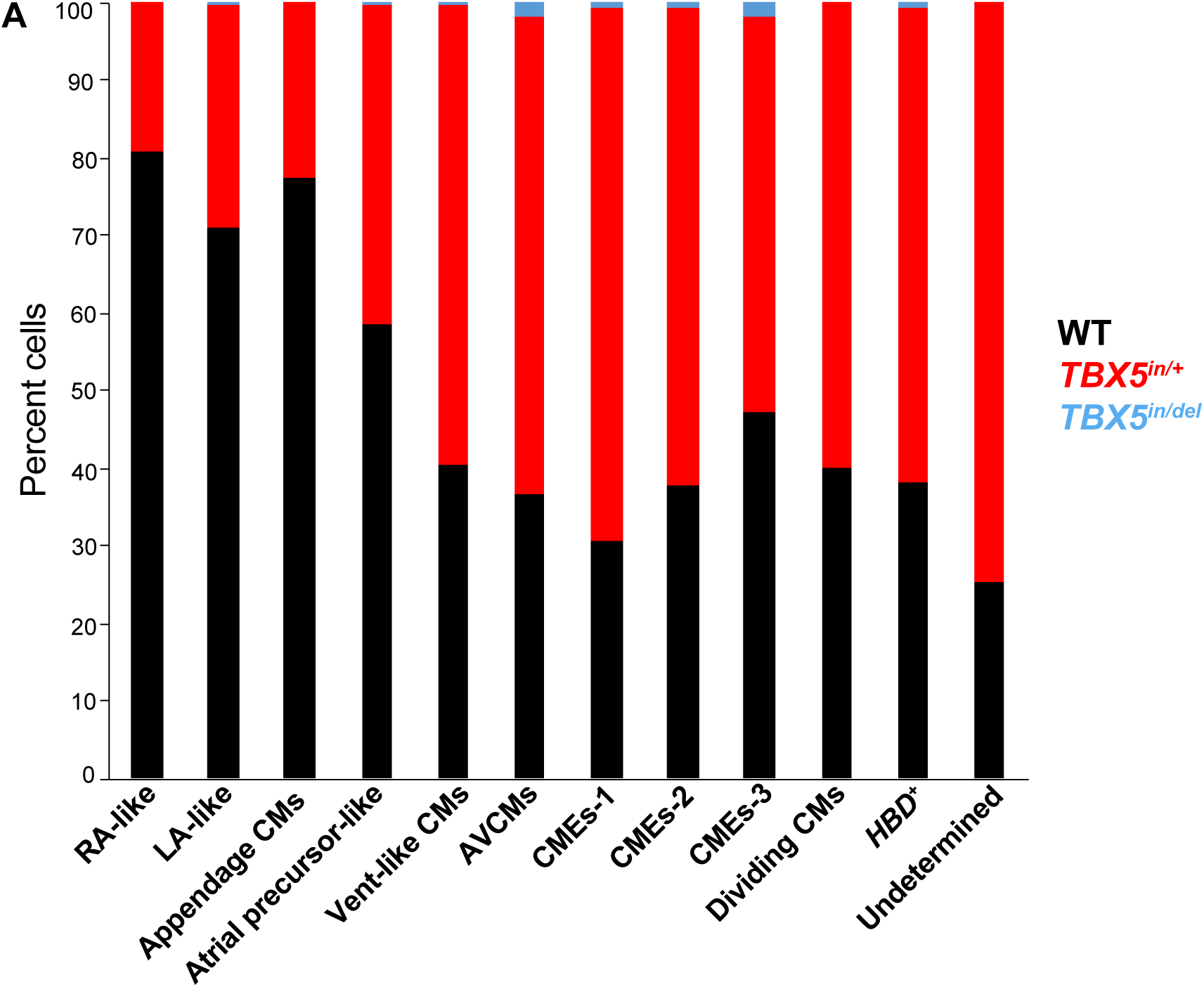
Analysis of CMs in a human cellular model of atrial disease. (A) Bar plot shows CM subtypes at day 20 by *TBX5* genotypes. scRNA-seq of two biological replicates by *TBX5* genotype were analyzed.

**Fig. S6.**
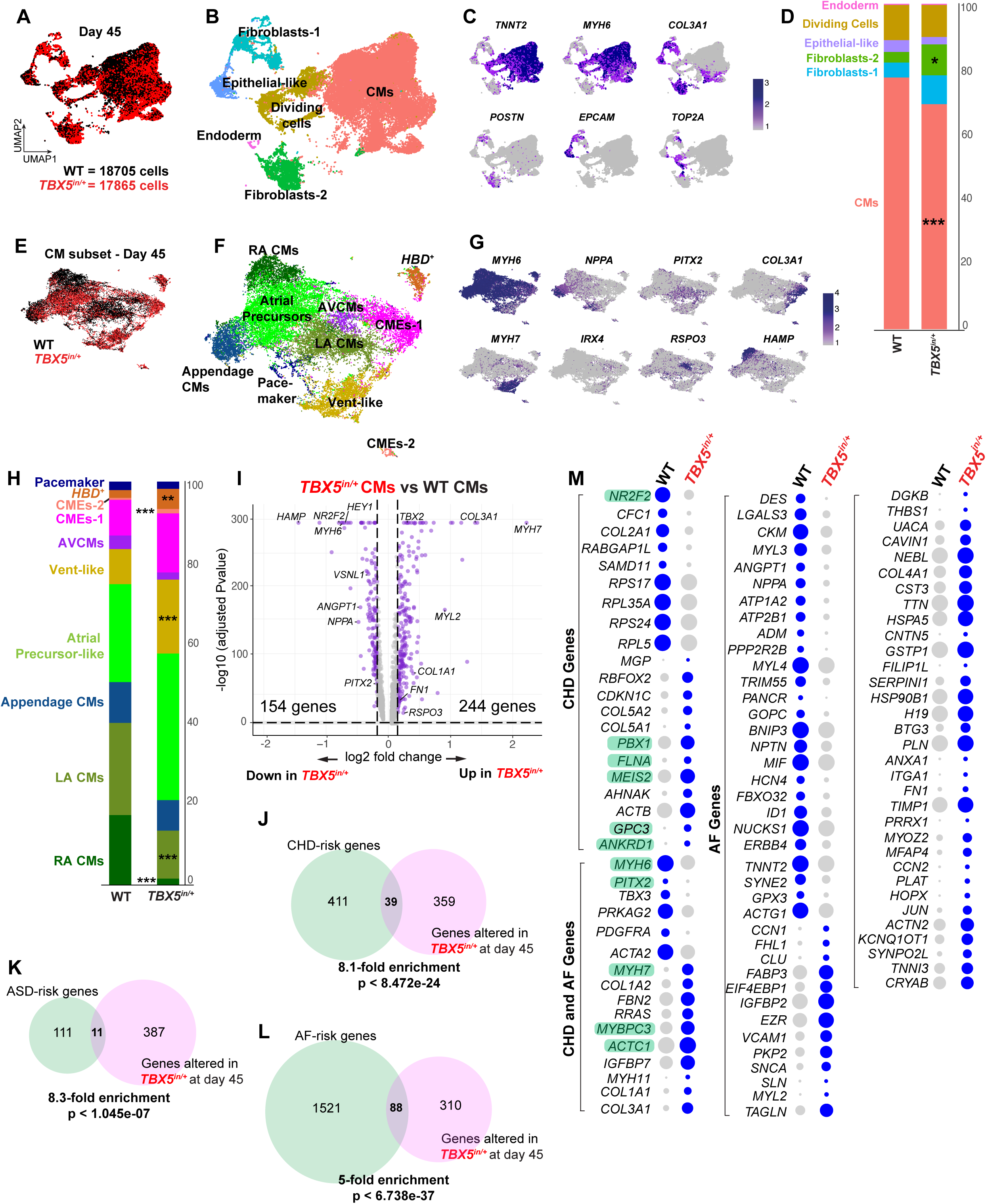
Dysregulated gene expression in human iPSC-derived atrial cells of *TBX5* heterozygous mutants at day 45 by scRNA-seq. UMAP by (A) genotype or (B) cell type label at day 45. (C) Feature plots of marker genes for cell type labels. (D) Bar plot shows distribution of cell types in WT or *TBX5^in/+^,* including a reduction in CMs in *TBX5* mutants (*adj. p<0.05, ***adj. p<0.001 by linear regression). UMAP of CM subtypes by (E) genotype or (F) CM subtype labels. (G) Feature plot of marker genes for CM subtype labels. (H) Bar plot shows distribution of CM subtypes in WT or *TBX5^in/+^,* including alterations to several CM subtypes (**adj. p<0.01, ***adj. p<0.001 by linear regression). (I) Volcano plot shows diferentially expressed genes (adj. p-value by Wilcoxan rank sum). (J-L) CHD-, ASD– and AF-risk genes are enriched among differentially expressed TBX5-sensitive genes at day 45 (non-adjusted p-value by hypergeometric test). Dot plots display differentially-expressed CHD, ASD (in green) and AF genes in (M) *TBX5^in/+^* (adj. p<0.05 by Wilcoxan rank sum). scRNA-seq of two biological replicates by *TBX5* genotype were analyzed.

**Fig. S7.**
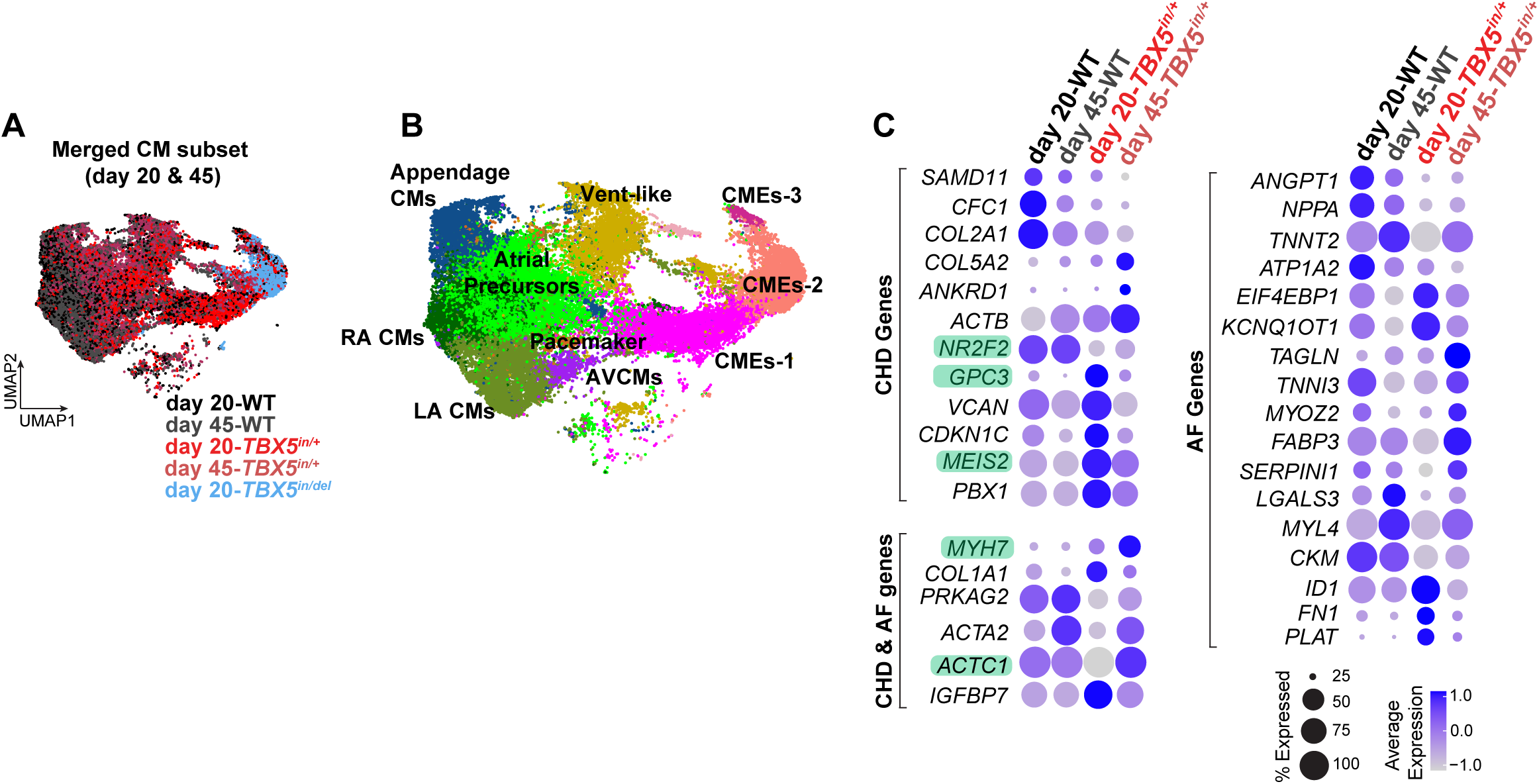
Comparison of differential gene expression in human iPSC-derived atrial cells of *TBX5* heterozygous mutants at day 20 and day 45. UMAP of CM subtypes in merged data of day 20 and day 45 by (A) *TBX5* genotype and (B) CM subtype labels. (C) Dot plots of differentially-expressed CHD, ASD (highlighted in green) and AF genes in *TBX5^in/+^* in merged data of day 20 and day 45 (adj. p-value<0.05 by Wilcoxan rank sum). scRNA-seq of two biological replicates by *TBX5* genotype at day 20 or day 45 were analyzed.

**Fig. S8.**
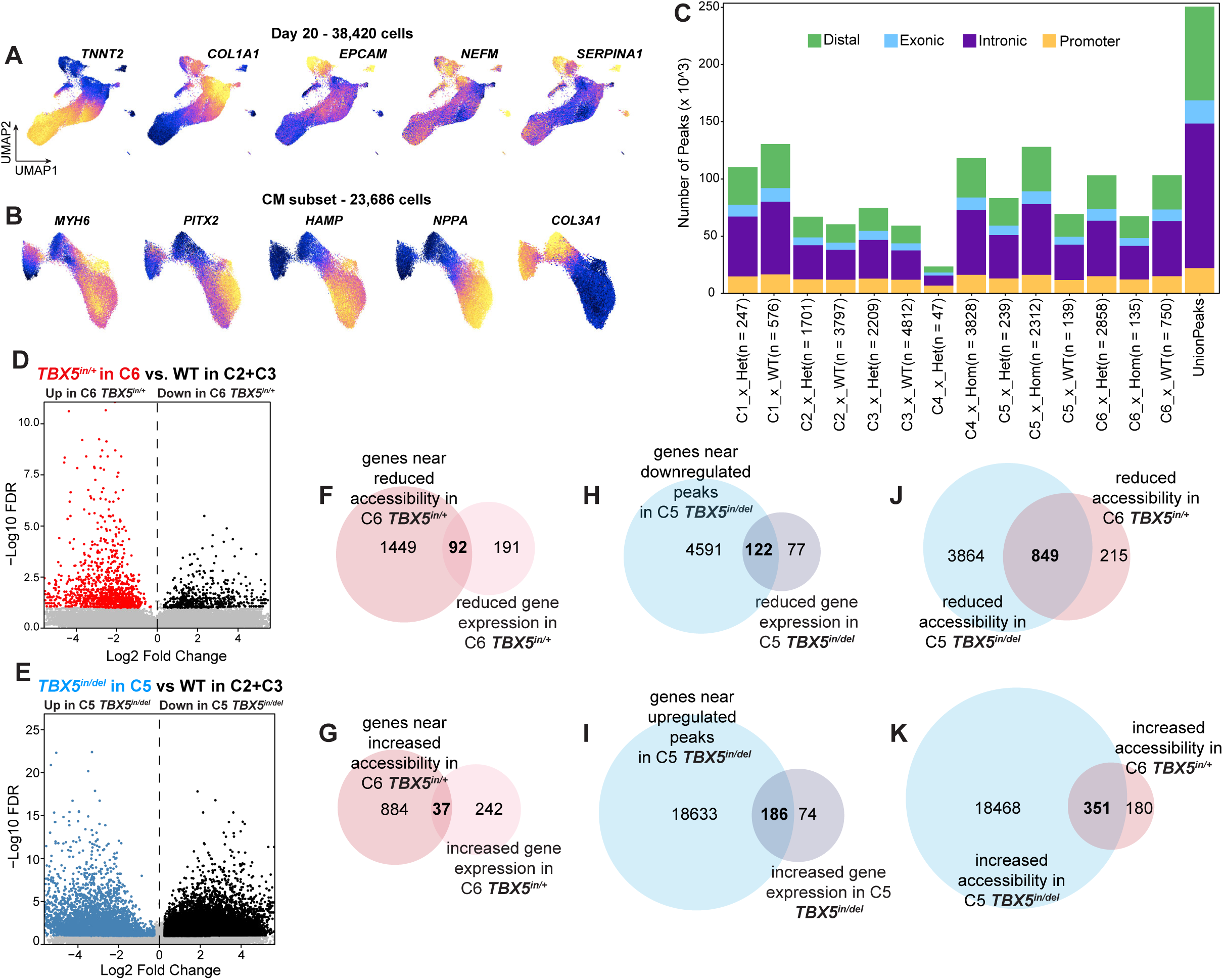
Analysis of chromatin accessibility in human iPSC-derived atrial cells of *TBX5* mutants. (A) Feature plots of gene scores identify putative cell types from chromatin accessibility. (B) Feature plots of gene scores identify putative CM subtypes from chromatin accessibility. (C) Genomic distribution of peaks by cluster. Differentially accessible peaks in (D) *TBX5^in/+^*or (E) *TBX5^in/del^*. (F-I) Overlap of genes near DARs and differential gene expression in *TBX5* mutants. (J, K) Overlap of DARs in *TBX5^in^*^/+^ and *TBX5^in^*^/*del*^. Two biological replicates of snATAC-seq by *TBX5* genotype were analyzed.

**Fig. S9.**
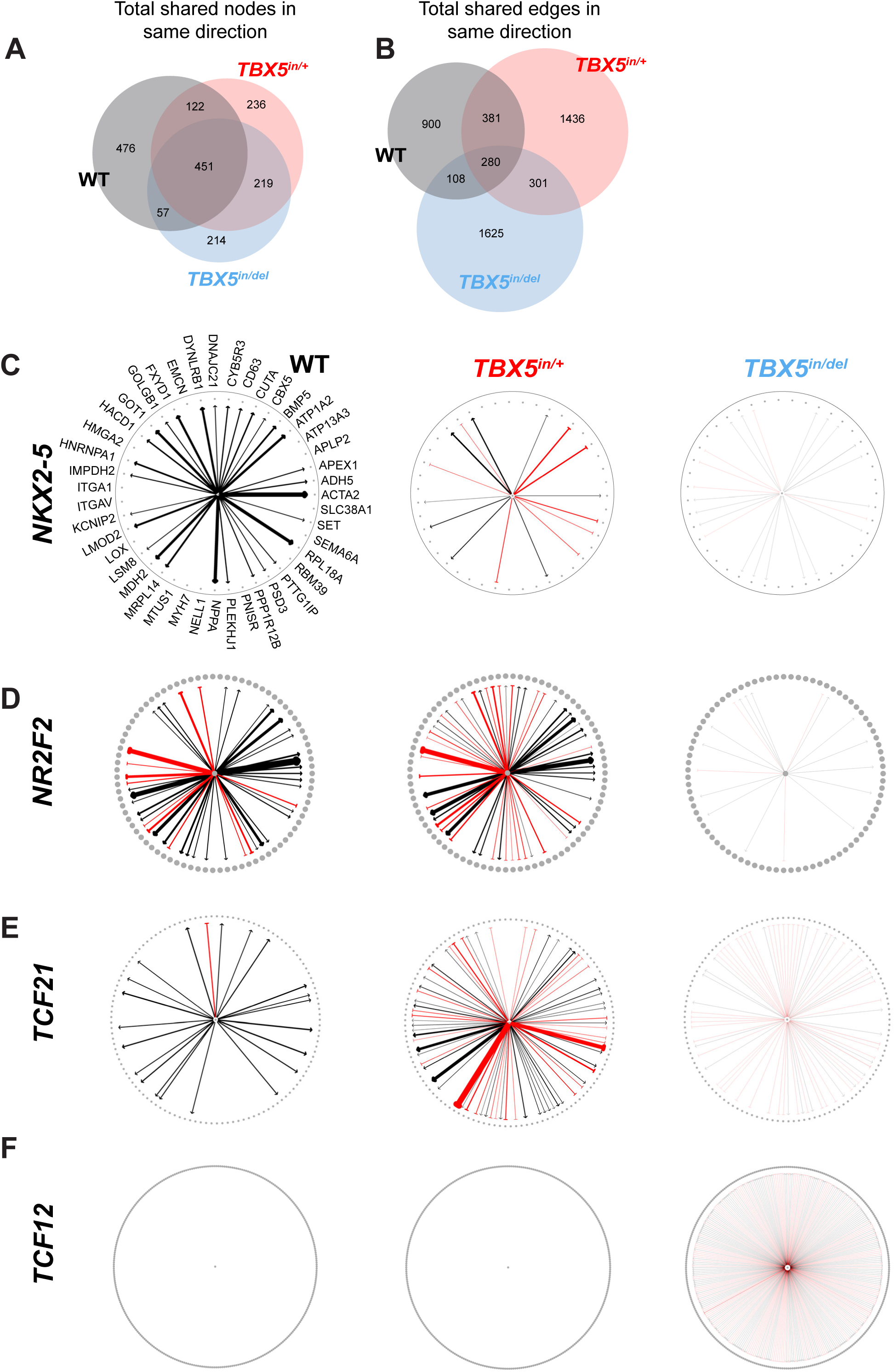
Network analysis of *TBX5* mutants in human iPSC-derived atrial CMs. Venn diagrams show overlap of (A) nodes or (B) edges in the same direction in *TBX5* mutants. Wheel diagrams of (E) ASD– and AF-risk *NKX2-5*, (F) ASD-risk *NR2F2*, (G) fibroblast-associated *TCF21*, and (H) fibroblast-associated *TCF12*. Strength of black arrows depicts strength of positive correlation, while strength of red arrows depicts strength of negative correlations. GRNs were derived from scRNA-seq or snATAC-seq of two biological replicates by *TBX5* genotype.

**Fig. S10.**
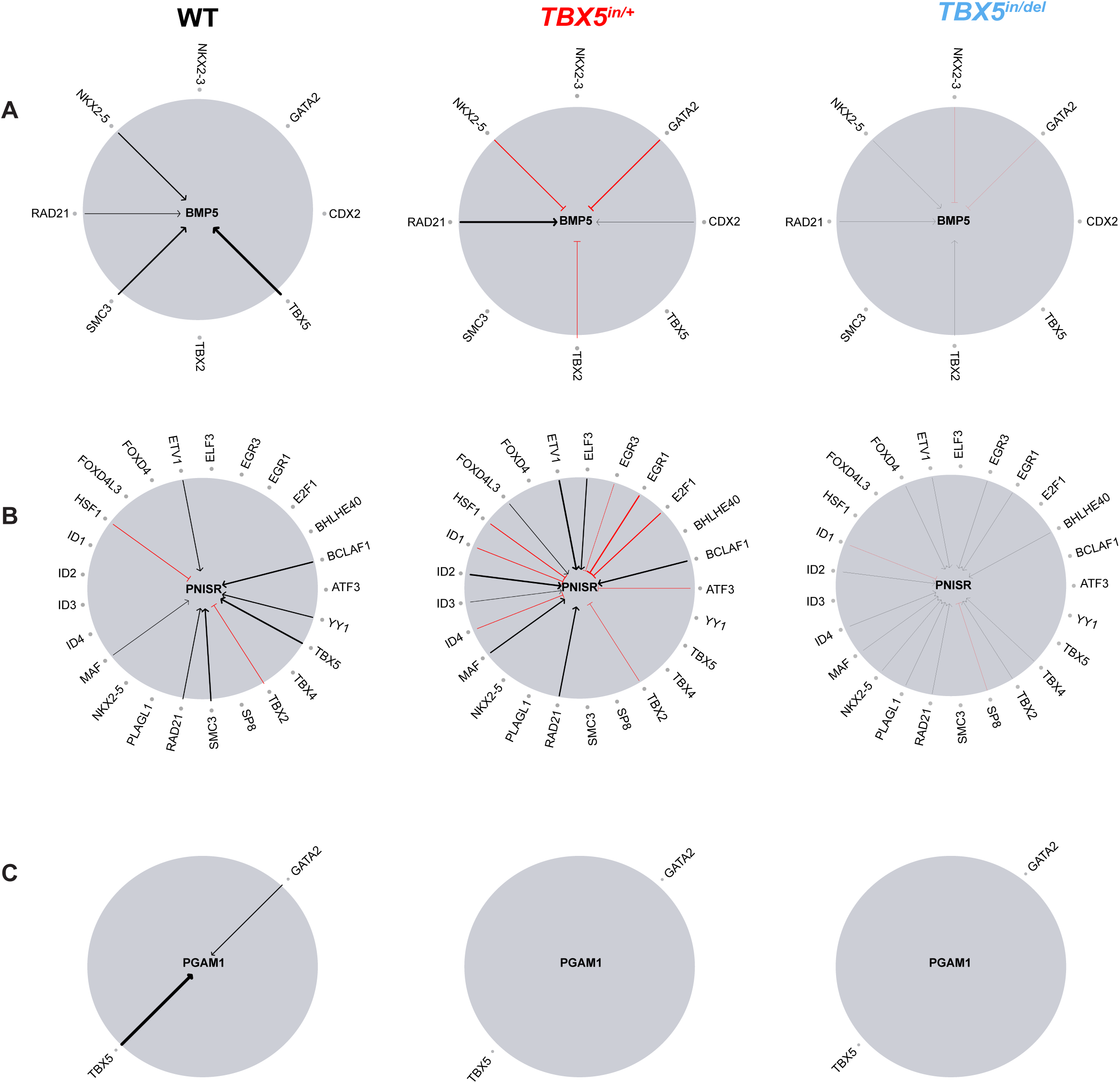
Altered connections to TBX5-dependent target edges in human iPSC-derived atrial CMs of *TBX5* mutants. Inverse wheel diagrams show connections of (A) *BMP5*, (B) *PNISR* and (C) *PGAM1* to other TFs that are gained or lost from reduced TBX5 dosage. GRNs were derived from scRNA-seq or snATAC-seq of two biological replicates by *TBX5* genotype.

**Table S1.** Table of differentially expressed genes between an *MYH6*^+^-enriched cluster of atrial CMs and an *MYL2*^+^-enriched cluster of ventricular CMs.

**Table S2.** Table of differentially accessible regions between *MYH6*-gene score^+^ atrial CM and *MYL2*-gene score^+^ ventricular CM clusters.

**Table S3.** Table of sources and targets in networks for atrial CM or ventricular CM clusters.

**Table S4.** Table of differentially expressed genes among *TNNT2*^+^ clusters by TBX5 genotype.

**Table S5.** Lists of disease-risk genes for CHDs, ASDs, AF, and ASDs and AF.

**Table S6.** Table of differentially accessible regions of *TNNT2*-gene score^+^ CM clusters by *TBX5* genotype.

**Table S7.** Table of sources and targets in networks for CM clusters from atrial CM differentiation by *TBX5* genotype.

